# Mitochondrial dysfunction plays a key role in premature musculoskeletal aging in hypochondroplasia mouse model

**DOI:** 10.64898/2026.10.16.759608

**Authors:** Chantal Fayad, Giulio Gatto, Stéphane Blouin, Michal Tamáš, Markus A. Hartmann, Adriana Figueroa-Garcia, Dimitrije Milunov, Sara Ceccacci, Ines Metatla, Ida Chiara Guerrera, Soham Saha, Paolo Bonaldo, Frank Zaucke, Nadja Fratzl-Zelman, Laurence Legeai-Mallet

## Abstract

Hypochondroplasia (HCH) is a developmental disorder characterized by short stature and caused by gain-of-function mutations in fibroblast growth factor receptor 3 (FGFR3). These mutations result in impaired bone and cartilage growth, tissues which are significantly affected in aging. We show that *Hch* mice exhibit hallmarks of accelerated aging: bone fragility, articular cartilage and subchondral bone changes, decreased muscle strength and reduction in muscle fibers size. Proteomic analyses of the cortical bone revealed that *Hch* mice follow a different aging trajectory than controls. The FGFR3 mutation leads to a premature boost of mitochondrial processes forcing mutants to reach an aged state much earlier than controls. We highlight a key role of mitochondria in the degradation of cortical bone of *Hch* mice, providing insight into mechanisms underlying premature aging in HCH, and offering new potential pathways involving mitochondria and FGFR3 to study aging.

## Introduction

Hypochondroplasia (HCH, OMIM #146000) is an autosomal dominant skeletal dysplasia characterized by rhizomelic dwarfism. Its clinical features include a stocky build, short hands and feet with brachydactyly, mild joint laxity, and scoliosis. Central hypotonia may be observed, and patients are at elevated risk of musculoskeletal complication such as osteoarthritis (OA). HCH is caused by gain-of-function mutations in fibroblast growth factor receptor 3 (FGFR3), ^1^. The FGFR3 variant, pAsn540Lys, located in the TK (tyrosine kinase) domain I of the protein, is responsible for 70% of cases of HCH. The estimated rate of occurrence of HCH is 1 in 15,000 to 40,000 newborns around the world ^2,3^. FGFR3 is a negative regulator of endochondral bone growth and variants cause a higher activation of the receptor and its downstream signaling pathways ^4,5^.

To study the precise pathophysiology of HCH, Loisay et al generated a mouse model (*Fgfr3^Asn534Lys^*^/+^) carrying the murine equivalent of the most frequent human HCH mutation, (p.Asn540Lys) ^6^. Cartilage and bone analyses have revealed the detrimental effect of a gain-of-function mutation in the FGFR3 gene on the formation of long bones during growth ^6^. Disease manifestations shift and transform across the continuum of adulthood and aging. Indeed, aging is a continuous process of functional loss, cumulative decline in the physiological function of an organism that has significant implications on human health and disease. The FGFR family and their ligands fibroblast growth factor (FGF) family have been shown to play a central role in the aging of skeletal tissues, regulating osteoblast function and mineral metabolism in the adult skeleton ^7^. For instance, in postmenopausal osteoporosis mouse models, recombinant FGF18 prevented bone loss by enhancing osteoblastogenesis and protecting osteoblasts from oxidative stress-induced apoptosis via Bcl-2 upregulation ^8^. FGF signaling has also emerged as a central player in onset and progression of OA. FGF7 acts as a catabolic factor in the joint, promoting cartilage destruction, subchondral bone thickening, and premature growth plate closure in mice whereas Fgf7 knockout mice exhibited significantly increased growth plate thickness (Shin et al., 2025). FGF18 acts through FGFR3 and suppresses chondrocyte hypertrophy in mice ^11^ and an intra-articular injection of FGF18 decreased cartilage degradation and matrix metallopeptidase 13 expression in rat post-traumatic OA^12^. A receptor-level shift occurs with aging: FGFR3 and FGF18 dominate in young mandibular cartilage of mice, sustaining homeostasis, whereas aging drives a progressive rise in FGFR1 expression and a concurrent decline in FGFR3, tipping the balance irreversibly toward catabolism and OA ^13^.

FGFs also play a role in the skeletal muscle. FGF2 reduces the loss and injury of skeletal muscle by activating the downstream RAS-MAPK signaling cascade via FGFR1. Notably, FGF expression declines progressively with age, contributing to age-associated reduction in muscle growth ^14^. According to a cross-sectional study, a cohort of older adults with sarcopenia had significantly higher serum FGF21 levels and significantly lower serum FGF19 levels than people without sarcopenia ^15^.

In this study, we distinguished between pathological changes and the normal aging process, with the goal of identifying the mechanisms underlying HCH-related aging. To address this issue, we studied the effect of the FGFR3 variant on the musculoskeletal system in the *Fgfr3^N534K/+^* (*Hch*) mouse model during aging ^6^. Our data revealed that bone metabolism is affected leading to an overall increase in tissue mineralization with age, whereas the mutation impacted size and number of osteocyte lacunae resulting in bone fragility. We discovered that *Hch* mice presented OA-like changes in articular cartilage, an increase in subchondral bone remodeling and sclerosis and structural and functional alteration of the muscle. Moreover, comparative proteomic analyses of cortical bone from mice of all ages highlighted that the *Hch* mutation leads to a premature acceleration of metabolic processes, particularly those related to mitochondria, such as mitochondrial aging and oxidative stress. Taken together, our findings provide a better understanding of the aging of the musculoskeletal system in HCH and highlight the key role of specific mitochondrial-related proteins and pathways in cortical bone.

## Results

### Bone metabolism is affected in *Hch* mice during aging

To determine the specific effect of the *Fgfr3* variant (*Asn534Lys*) on the appendicular skeleton during aging, we performed macroscopic and μCT analyses of femurs of 12 and 24-month-old male *Fgfr3^+/+^*(WT) and *Fgfr3^Asn534Lys/+^*(*Hch*) mice. As expected, femur length was significantly reduced in *Hch* mice compared to WT mice at 12 and 24 months of age (Extended Fig. 1a and Supplementary Table 1). In the trabecular bone of the WT femurs, we observed a significant decrease of bone mineral density (BMD) and bone volume/tissue volume ratio (BV/TV) between 12 and 24 months, consistent with normal aging (Extended Fig 1b,c). Comparative analyses of *Hch* and WT long bones revealed a significant decrease in the BV/TV ratio and BMD in the trabecular bone of the *Hch* mice regardless of their age (Extended Fig 1b,c and Supplementary Table 2).

Consistent with the aging process, cortical bone parameters of femoral mid-shafts showed a significant decrease in cortical BV/TV in WT animals (Fig. 1a,b). Interestingly, cortical bone analyses showed a significant increase in tissue mineral density (TMD) and BV/TV at 12 months of age in *Hch* mice compared to WT mice (Fig. 1b and Supplementary Table 3). These results were consistent with a previous study conducted in younger *Hch* mice ^6^. Altogether, these data demonstrated that regardless of the aging process, activation of the *Fgfr3* signaling pathway led to a decreased BMD in trabecular bone and an increased TMD in cortical bone. These results highlight a direct impact of the Fgfr3 variant on bone metabolism leading to an increased risk of associated bone fragility.

**Figure 1:**
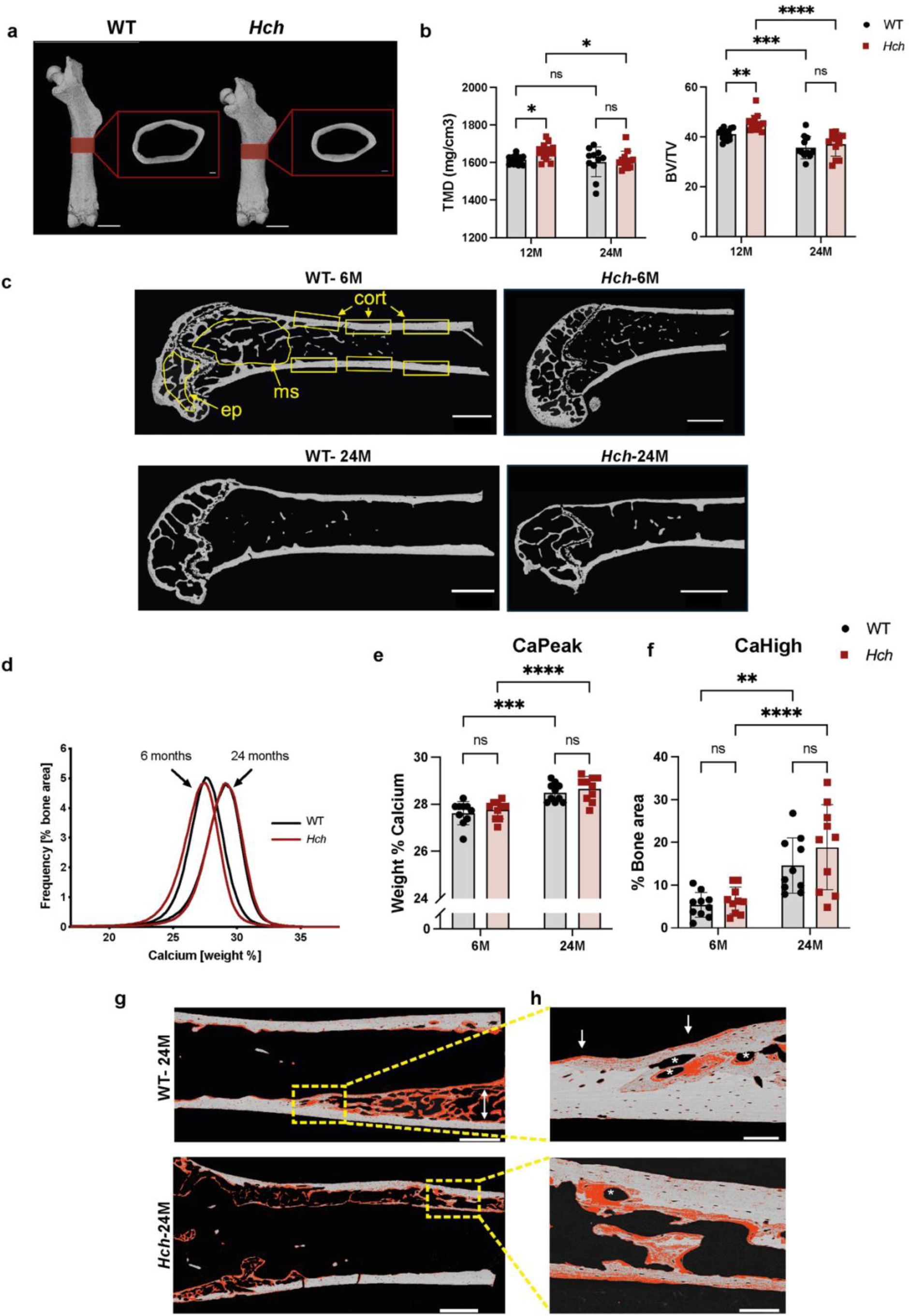
Bone metabolism is affected in *Hch* mice during aging. **a**) 3D visualization of representative 12-month-old WT and *Hch* male femurs. Highlighted in red indicates the ROI chosen for the cortical bone studies. Scale bars: 500 μm for the whole bone and Scale bars: 200 μm for cortical parts. (**b**) Histograms for the cortical bone parameters of 12 & 24-month-old femurs of *Hch* mice (n=15, n=11 respectively) vs WT mice (n=14, n=12 respectively): tissue mineral density (TMD) and Bone volume/total volume (BV/TV). (**c**) Representative overview images obtained by quantitative backscattered electron imaging (qBEI) of longitudinal sections of murine proximal femora of 6-month-old and 24-month-old WT and *Hch* male mice showing the three regions of interest that were evaluated separately: metaphyseal spongiosa (ms), epiphysis (ep), and cortical bone (cort). The six rectangles in the cortices indicate the six images used for the analysis of osteocyte lacunae sections (OLS) later. Scale bars: 1 mm (**d**) Bone mineralization density distribution (BMDD) histograms of cortical bone (cort) from 6 month and 24-month-old samples, obtained by quantitative backscattered electron imaging (qBEI). **(e,f)** Statistical comparison of CaPeak and CaHigh of *Hch* mice (n=10) vs WT mice (n=10) in the cortex. CaPeak: the most frequently measured calcium concentration. CaHigh: the percentage of highly mineralized bone area (**g**) Cortical bone area from a 24-month-old WT and *Hch* mice showing intense new bone formation. Red overlay indicates areas mineralized below 25.3 weight % calcium. The outer compact cortices are not highlighted in red, mirroring a higher mineral content and therefore older tissue age, whereas the less mineralized bone tissue marked by the white arrow is highly porous. (**h**) Higher-magnification detail of (**g**) showing the transition area from compact cortical bone to a zone of intracortical bone remodeling. White asterisks indicate osteon-like structures with concentric new bone formation, typical of intracortical bone remodeling. White arrows point to endosteal regions stained in red, indicating de novo bone apposition. Note that these highly porous cortical areas were not included in the cortical BMDD values. Data are presented as mean ± SD. (**\*\*\*\*,** *P* < 0.0001; \*\*\**, P* < 0.001; \*\**, P* < 0.01; \**, P* < 0.05; ns, non-significant; Mann-Whitney for 2 groups and two-way ANOVA for >2 groups).

We further evaluated bone tissue characteristics by quantitative backscattered electron imaging (qBEI) in longitudinal sections of femoral bone from 6- and 24-month-old male mice to assess bone matrix mineralization in the metaphysis, epiphysis, and cortex—three key skeletal sites that differ in tissue age (Fig. 1c). In accordance with previous studies ^16–18^, we observed a progressive increase from the metaphyseal spongiosa to the epiphysis to cortical bone in CaMean and CaPeak in 6-month-old animals for both genotypes (Extended Fig. 2a,b, Supplementary Table 4). We observed no differences between WT and *Hch* mice in any of the bone mineralization density distributions (BMDD) at any of the bone sites (Supplementary Table 5,6 and 7). The BMDD of cortical bone obtained by qBEI showed that the peak positions of the WT and *Hch* samples do not differ at either 6 or 24 months of age. The BMDD curves of the 24-month-old mice are shifted to the right, toward higher mineral content, indicating that in both genotypes the degree of mineralization increases with age (Fig. 1d). qBEI analyses of the cortical bone showed that CaMean, CaPeak and Ca High were not different between genotypes but they increase with age in both genotypes while CaLow was unchanged (Fig. 1e,f, Extended Fig. 2c,d and Supplementary Tables 5,6 and 7). While analyzing the samples, we unexpectedly observed intense new bone formation in the cortical bone area from some 24-month-old WT and *Hch* mice (Fig. 1g). Higher-magnification detail shows the transition area from compact cortical bone to a zone of intracortical bone remodeling (Fig. 1h). Altogether, these analyses indicate that aging modify bone metabolism, in both genotypes.

### Osteocyte lacunae sections (OLS) density is higher in *Hch* male mice

Having established that bone metabolism was modified by aging, we next addressed the impact of Fgfr3 itself on bone cells. Osteocyte lacunae sections (OLS) were evaluated in the longitudinal sections of cortical bone ^19^ (Fig.1c and Fig. 2a). Analyses of *Hch* versus WT femoral cortex reveals elevated OLS density (Fig. 2b) and, in accordance, a decrease in OLS-NND (nearest neighbor distance) in *Hch* mice at 6 months of age (Fig. 2c). OLS area and perimeter decreased with age in both genotypes and were smaller in *Hch* mice than in WT at 6 months of age (Fig. 2d, Extended Fig. 2e and Supplementary Table 8). No differences in OLS size were found between genotypes at 24 months of age (Fig. 2d, Extended Fig. 2e and Supplementary Table).

**Figure 2:**
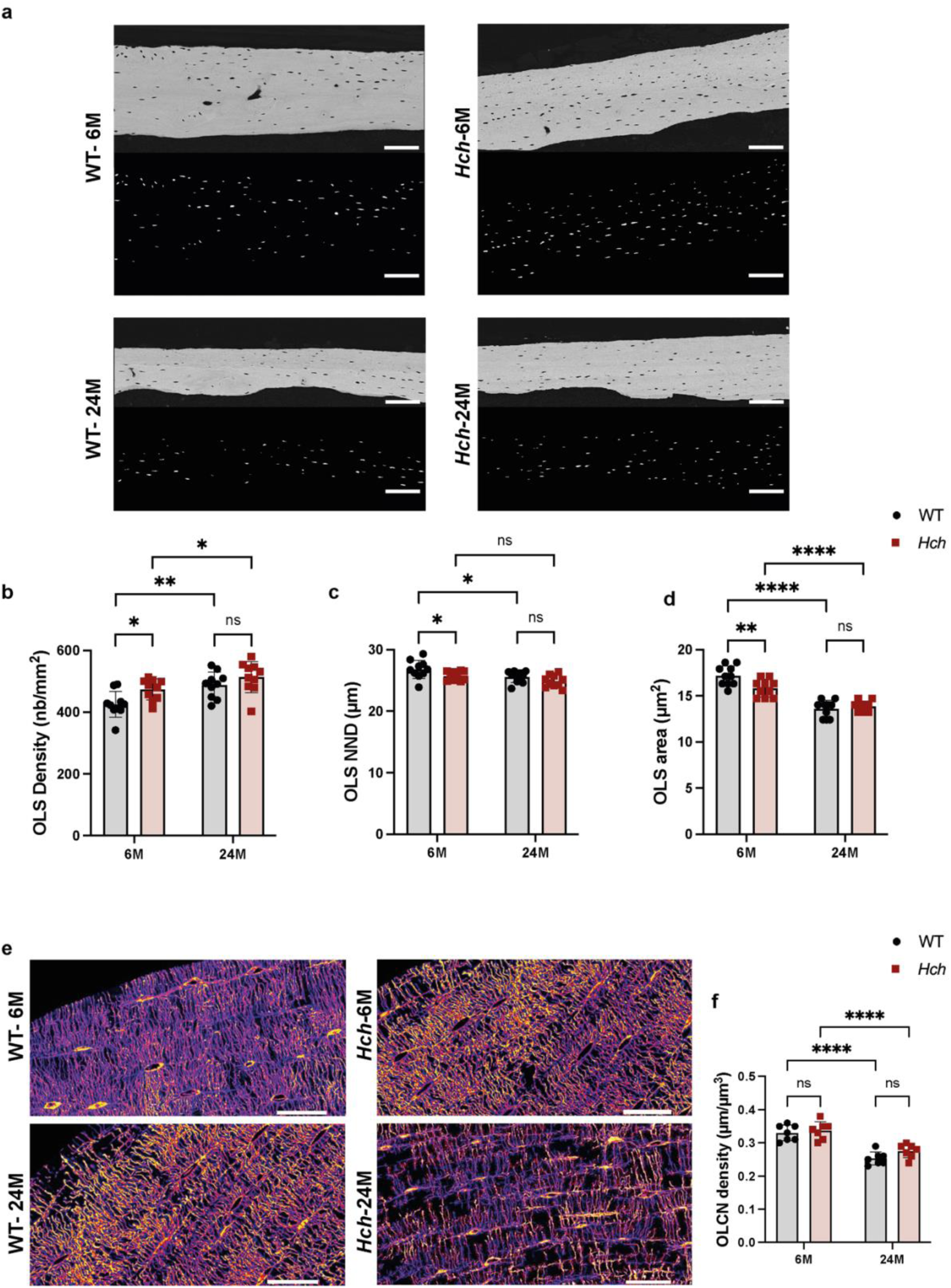
*Hch* mutation affects osteocytes lacunar and canalicular characteristics. **(a)** Representative qBEI images of the cortex and corresponding binary images showing the osteocyte lacunae sections (OLS) included in the quantification for each group. Scale bars: 100 µm. (**b,c,d**) Statistical comparison of OLS variables in *Hch* (n = 10) versus WT mice (n = 10) mice femoral cortex. (**e**)Representative confocal laser scanning microscopy (CLSM) images of rhodamine-stained cortical bone from 6-month-old and a 24-month-old WT and *Hch* mice showing the osteocyte lacuno-canalicular network (OLCN). Scale bar: 25 µm. (f) Statistical comparison of OLCN in *Hch* (n = 10) versus WT mice (n = 10) mice. Data are presented as mean ± SD. (**\*\*\*\****, P* < 0.0001; \*\*\**, P* < 0.001; \*\**, P* < 0.01; \**, P* < 0.05; ns, non-significant; Mann-Whitney for 2 groups and two-way ANOVA for >2 groups).

### Osteocyte lacuno-canalicular density decreases with age in both genotypes

We performed rhodamine-staining in cortical bone for WT and *Hch* (Fig. 2e) and evaluated osteocyte lacuno-canalicular network (OLCN) characteristics in cross-sectioned femoral bone by confocal-laser scanning microscopy imaging ^18,20^. Analyses revealed a decrease of OLCN density with age but no differences between genotypes (Fig. 2f and Supplementary Table 9). Accordingly, the OLCN appears denser in bone tissue from the younger compared to the older mice (Fig. 2e) indicating a deterioration of osteocyte connectivity and communication within the bone matrix with aging.

### Spinal canal stenosis is observed in *Hch* mice

To investigate both the impact of Fgfr3 variant and the effect of bone aging on the axial skeleton, we investigated bone microarchitecture in vertebrae. First, we examined the shape of the fifth lumbar (L5) vertebrae and found that *Hch* mice showed significant reduction in vertebral body length at 12 and 24 months of age, respectively, as well as a reduction in interpedicular distance at 12 months of age (Fig. 3a,b,c and Supplementary Table 10). Analyses of the vertebral canal revealed a significant reduction of its surface area in *Hch* mice compared to WT littermates at 12 and 24 months of age (Fig. 3d,e and Supplementary Table 10). These observations support the idea that the Fgfr3 gain-of-function variant causes the canal vertebral stenosis as reported in human ^21,22^.

**Figure 3:**
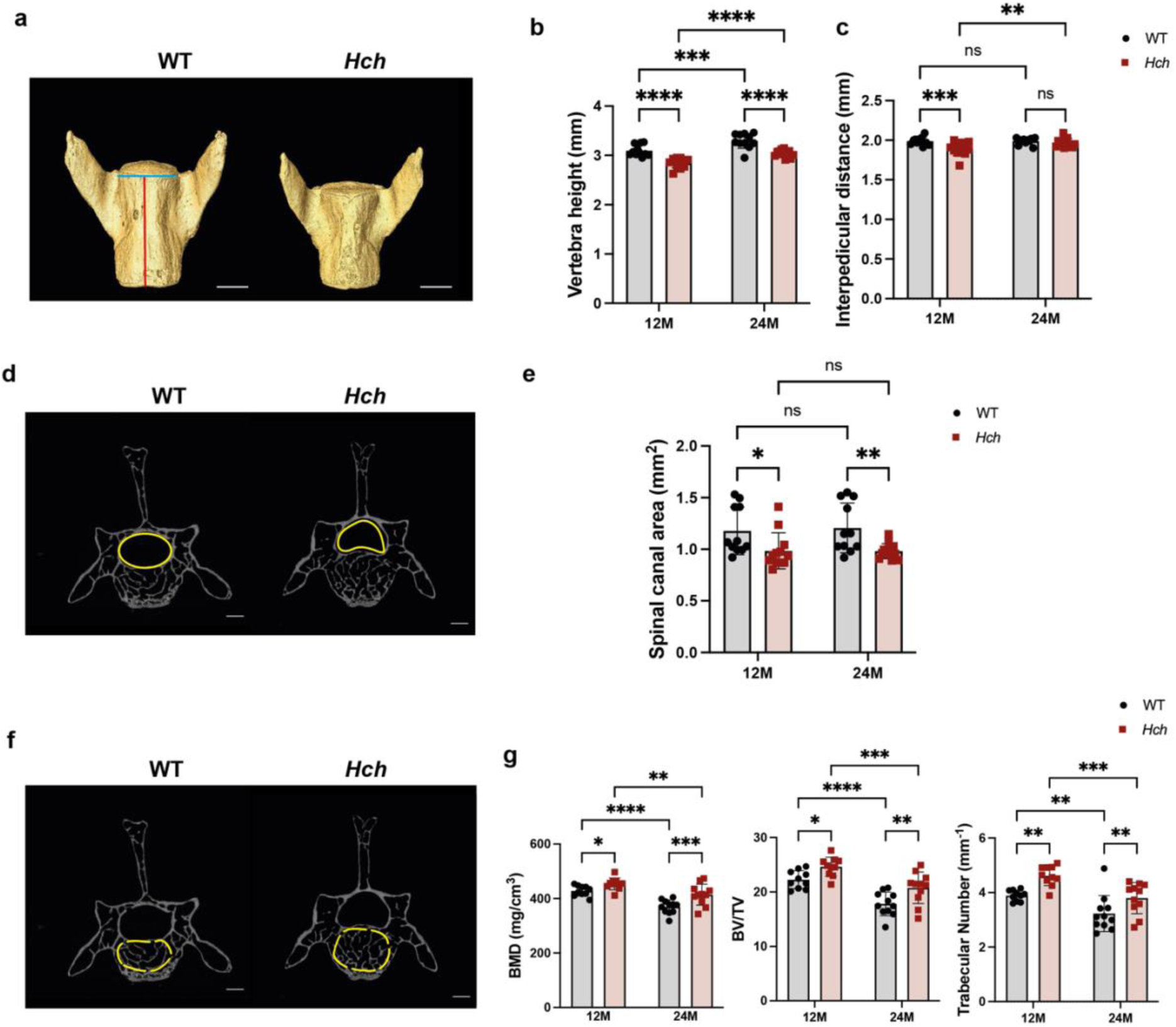
*Hch* mutation results in altered trabecular architecture and stenosis of spinal canal in vertebrae. (**a**) 3D rendering of representative 24-month-old WT and *Hch* male mice L5 vertebrae. Scale bar: 500 μm. (**b,c**) Graphical representations of L5 height and interpedicular distance of 12-month-old and 24-month-old WT (*n* = 11 for both) and *Hch* (*n* = 12 & n=13 respectively). The vertebrae height is highlighted in red, while the interpedicular distance is in blue. (**d**) 3D visualization of representative 24-month-old WT and *Hch* L5 trabecular bone. The spinal canal area is highlighted in yellow. Scale bar: 500 μm (**e**) Graphical representation of L5 spinal canal area of 12-month-old and 24-month-old WT *(n* = 11 for both) and *Hch* (*n* = 12 & n=13 respectively). (**f**) 3D visualization of representative 24-month-old WT and *Hch* L5 trabecular bone. The area of trabecular bone ROI is highlighted in yellow dashed line. Scale bar: 500 μm. (**g**) Histograms for the trabecular bone parameters of 12 & 24-month-old L5 obtained by μCT: Bone mineral density (BMD), bone volume/total volume (BV/TV) and trabecular number (Tb.N). Data are presented as mean ± SD. (**\*\*\*\*,** *P* < 0.0001; \*\*\**, P* < 0.001; \*\**, P* < 0.01; \**, P* < 0.05; ns, non-significant; Mann-Whitney for 2 groups and two-way ANOVA for >2 groups).

We then studied the bone aging process of the vertebral trabecular bone in WT mice aged 12 and 24 months. We observed a significant decrease of the BMD, BV/ TV and trabecular number which is consistent with our previous data on the appendicular skeleton (Extended Fig 1b,c). Next, we evaluated the same parameters for *Hch* mice. The analyses showed an increase in BMD, BV/TV and in trabecular number (Tb.Nb) in *Hch* compared to WT mice at 12 and 24 months of age (Fig. 3f,g and Supplementary Table 11). Together, these findings suggest that activation of the Fgfr3 gene induces a trabecular sclerosis phenotype in the vertebrae of *Hch* mice.

### The *Hch* variant promotes cartilage degeneration and increases subchondral bone formation during aging

We evaluated cartilage degeneration and subchondral bone (SB) changes specifically in the tibial compartment of the knee joint. The articular cartilage was analyzed in male mice at 12 and 24 months of age. Even though no differences in OARSI score ^23^ between both genotypes and between ages were observed (Extended Fig. 3a), the thickness of the articular cartilage in both the medial and lateral compartments was significantly decreased in *Hch* mice compared to WT at 12 and 24 months of age. The medial cartilage thickness further decreased significantly from 12 to 24 months though only in *Hch* mice (Fig. 4a,b and Supplementary Table 12). Evaluation of the calcified area in medial and lateral cartilage revealed differences only in the medial compartment. Here, the calcified area is significantly increased in *Hch* mice compared to WT at 12 months of age indicating an increased calcification caused by the *Hch* variant (Fig. 4a,c and Supplementary Table 12).

**Figure 4:**
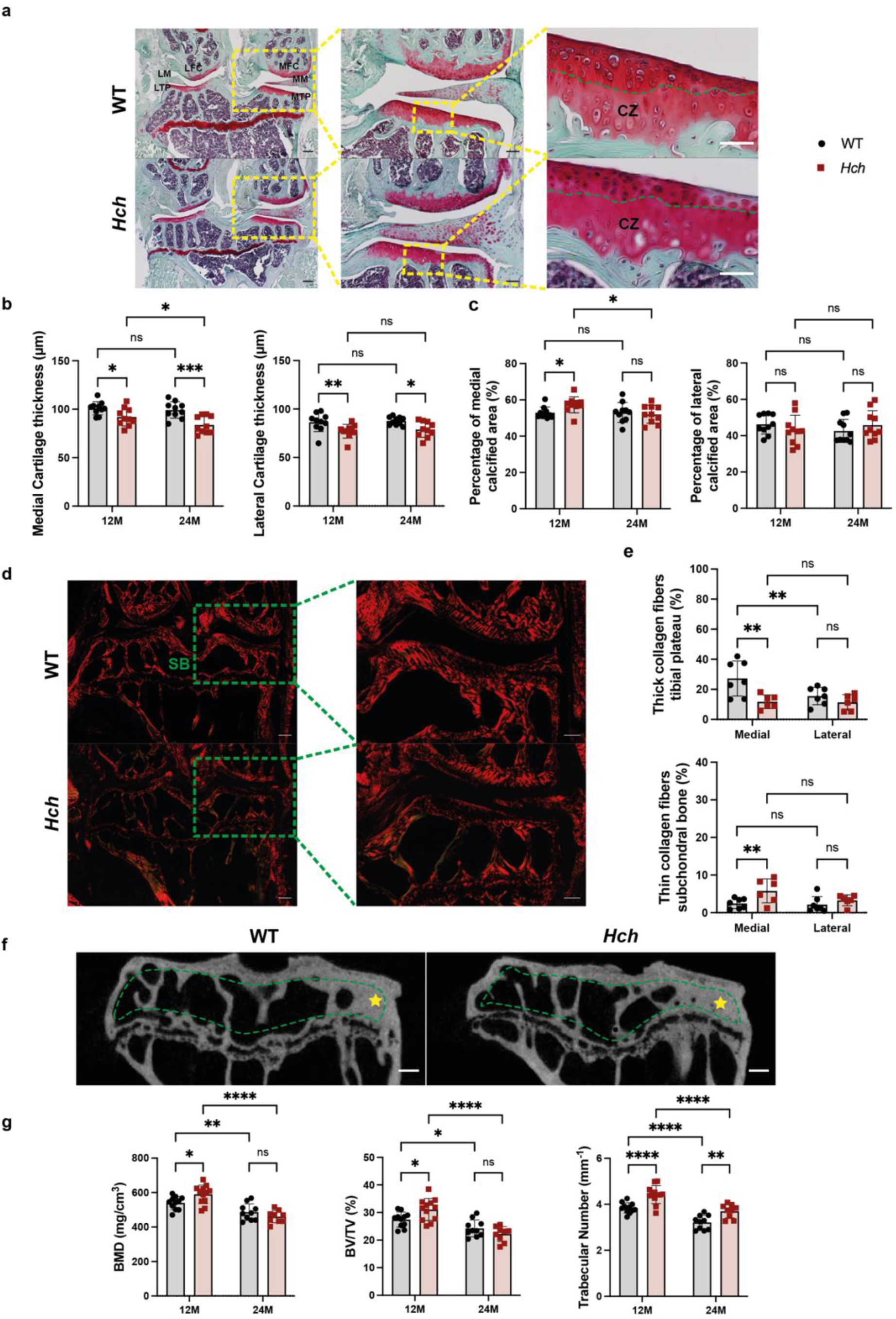
*Hch* mutation drives osteoarthritic cartilage changes coupled with subchondral bone remodeling during aging. (**a**) Representative safranin O/Fast green staining of frontal sections of right knee joints from 24-month-old male mice showing lateral and medial femoral condyles (LFC/MFC), lateral and medial tibial plateaus (LTP/MTP), lateral and medial menisci (LM/MM). Scale bar: 200 μm. Higher magnification of the MFC, MM and MTP. Scale bar: 100 μm. The hight magnification detail showing calcified zone (CZ), tidemark is represented in green dashed line Scale bars: 50 μm. (**b**) Graphical representation of medial and lateral cartilage thickness of 12-month-old (WT *n* = 10; *Hch n* = 10) and 24-month-old male mice (WT *n* = 10; *Hch n* = 10). (**c**) Percentage of calcified area in medial and lateral tibial cartilage of 12-month-old (WT *n* = 10; *Hch n* = 10) and 24-month-old male mice (WT *n* = 10; *Hch n* = 10). (**d**) Picrosirius red staining of frontal sections of right knee joints from 12-month-old male mice. Scale bars: 200 μm, Higher magnification of the subchondral bone (SB). Scale bars: 100 μm (**e**) Percentage of thick collagen fibers in LTP/MTP, and percentage of thin collagen fibers in subchondral bone of 12-month-old mice (WT *n* = 7; *Hch n* = 6). (**f**) Representative μCT images of SB in 12-month-old WT and *Hch* male mice. Sclerosis is represented with a yellow star and the analyzed area with a green dashed contour. Scale bars: 200 μm. (g) Histograms representation of bone mineral density (BMD), bone volume/total volume (BV/TV) and trabecular number (Tb.N) of SB of 12-month-old (WT *n* = 11; *Hch n* = 12) and 24-month-old (WT *n* = 10; *Hch n* = 9) male mice. Data are presented as mean ± SD. (**\*\*\*\*,** *P* < 0.0001; \*\*\**, P* < 0.001; \*\**, P* < 0.01; \**, P* < 0.05; ns, non-significant; Mann-Whitney for 2 groups and 2-way ANOVA for >2 groups).

In addition, to visualize collagen fiber in cartilage and subchondral bone, we analyzed and quantified picrosirius red staining in 12-month-old WT and *Hch* mice, using polarized light ^24^.We found a decrease of thick collagen fibers (red fibers) in the medial tibial plateau and an increase in thin collagen fibers (green fibers) in the medial part of the subchondral bone with no changes in the remaining parameters in *Hch* mice compared with WT (Fig. 4d,e, Extended Fig. 3b,c and Supplementary Table 13). Altogether, these data indicate that *Hch* mice exhibit early OA-like cartilage thinning with increased calcified cartilage area in the medial part of the tibial cartilage.

Analysis of the SB by μCT revealed a decrease in BMD, BV/TV and Tb. Nb between 12 and 24 months of age in WT animals indicating that aging affects the subchondral bone parameters. In contrast, compared to WT, *Hch* mice showed a significant increase in BMD and BV/TV at 12 months of age and an increase in the trabecular number at 12 and 24 months of age, indicating an increase in the subchondral bone remodeling (Fig. 4f,g). Interestingly, more subchondral sclerosis was observed in *Hch* mice, and again the medial compartment was more affected (Fig. 4f). Both genotypes showed a decline in BV/TV and BMD with age but this decline was significantly steeper in mutants, suggesting that the mutation accelerated age-dependent SB changes (Supplementary Table 14). Cartilage and subchondral bone changes were mainly seen in the medial compartment which may be attributable to different compartmental loading patterns. Here, we demonstrate that *Hch* mice exhibit phenotypic traits of OA in articular cartilage and subchondral bone that are accelerated by aging.

### *Hch* mice show compromised muscle homeostasis

Given the tight relationship between muscle and bone from embryogenesis through development and into aging, the changes in one tissue are likely to affect the other ^25^. We hypothesize that disruption of bone homeostasis in our *Hch* mouse model likely alters the osteokines that interact with the muscles, which would consequently lead to a muscular phenotype. Therefore, we examined the effect of *Hch* variant on skeletal muscles, and started by performing a grip strength test on 12-month-old male mice (Fig. 5a). Interestingly, *Hch* mice had reduced peak muscle force compared to WT at both ages (Fig. 5a and Supplementary Table 15). Given the involvement of bone-derived osteokines in controlling muscle homeostasis, we then investigated whether the atrophy program is changed in *Hch* muscle by measuring Trim63 (MuRF1) RNA expression level, a muscle-specific E3 ubiquitin ligase and canonical marker of muscle catabolism. Expression of the *Trim63* gene was significantly lower in tibialis anterior muscle from *Hch* mice compared to WT at 12 months of age (Fig. 5b), pointing to an altered protein turnover that could lead to impaired muscle homeostasis. The expression of Pax7, Myod1 and Myh3, markers of myogenesis, did not differ between WT and *Hch* muscle samples (Fig. 5b). We next assessed the muscle phenotype at the structural level by performing wheat germ agglutinin (WGA) surface staining on tibialis anterior muscle sections from 12-month-old mice (Fig. 5c). Cross sectional area and minimal Feret’s diameter were significantly decreased in *Hch* samples compared to WT (Fig. 5d-f), whereas the shape factor index was significantly increased (Fig. 5g), indicating smaller fibers and higher shape variability (Supplementary Table 16). Taken together, these data reveal the presence of structural and functional defects in skeletal muscles of *Hch* mice, suggesting impaired muscle homeostasis that may be influenced by changes described above in bone parameters in these mice.

**Figure 5:**
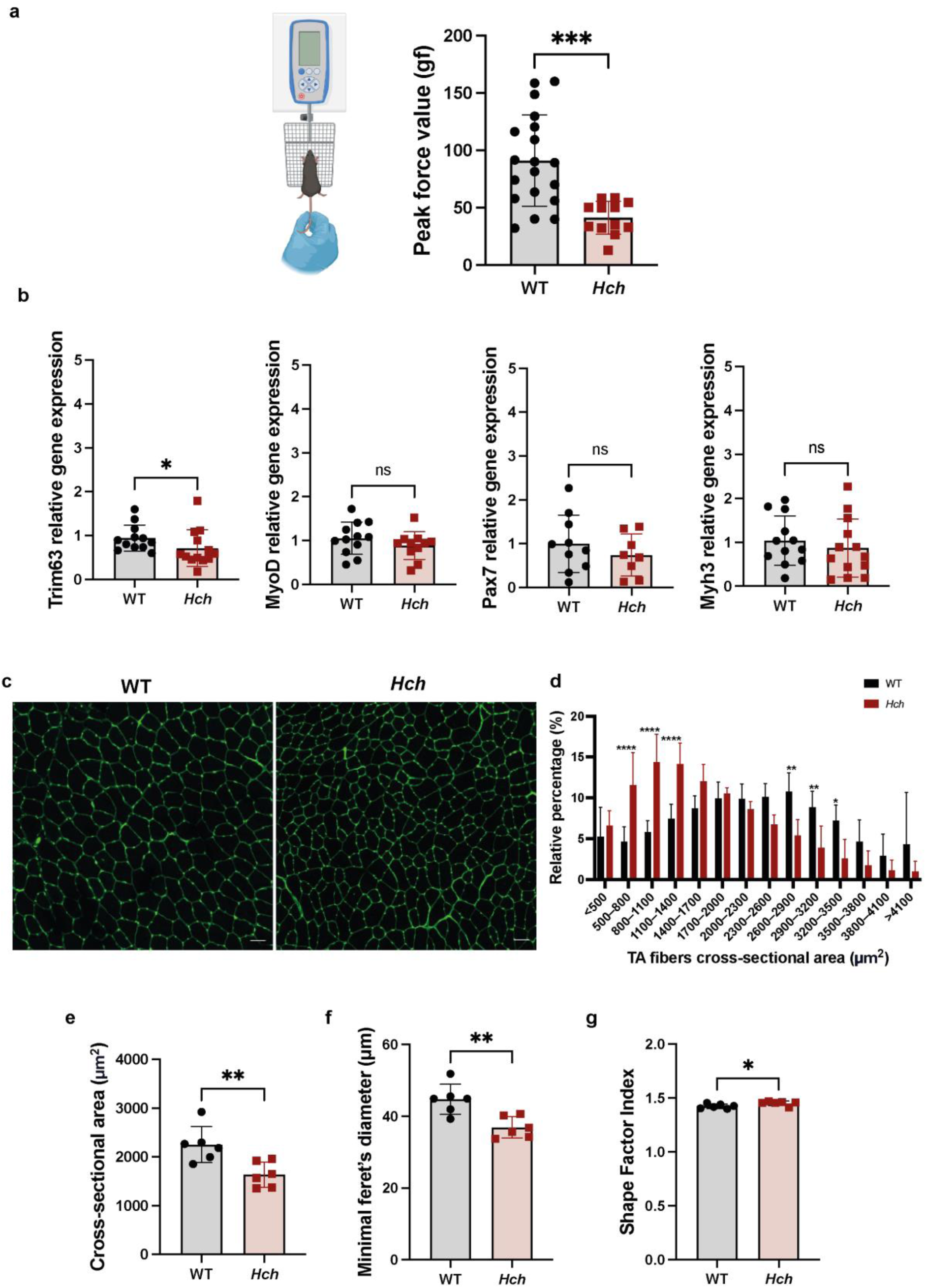
*Hch* mice showed reduced muscle’s strength and altered muscle’s function and shape. **(a)** Schematic diagram of grip strength test and histograms of peak force values of 12-month-old male mice (*WT n* = 19; *Hch n* = 12). (**b**) RT-qPCR analysis of tibialis anterior muscle samples from 12-month-old WT and *Hch* male mice. Trim63 relative gene expression in *WT* (*n* = 12) and *Hch* (*n* = 13), MyoD relative gene expression in *WT* (*n* = 12) and *Hch* (*n* = 11), Pax7 relative gene expression in *WT* (*n* = 10) and *Hch* (*n* = 8) and Myh3 relative gene expression in *WT* (*n* = 12) and *Hch* (*n* = 13) are shown. (c) Representative fluorescence images of WGA-stained tibialis anterior muscles from 12-month-old WT and *Hch* mice. Scale bar: 50 μm. (**d**) Distribution of myofibers with different cross-sectional area ranges in tibialis anterior muscles from WT (*n* = 6) and *Hch* (*n* = 6) mice. (**e-g**) Histograms representing mean cross-sectional area (**e**), minimal Feret’s diameter (**f**) and shape factor index (**g**) in tibialis anterior muscles from WT (*n* = 6) and *Hch* (*n* = 6) mice. Data are presented as mean ± SD. (**\*\*\*\*,** *P* < 0.0001; \*\*\**, P* < 0.001; \*\**, P* < 0.01; \**, P* < 0.05; ns, non-significant; Mann-Whitney for 2 groups and two-way ANOVA for >2 groups).

### Canonical aging and *Hch*-driven features rely on non-overlapping biological signaling

To elucidate the underlying mechanisms by which the gain-of-function variant of the *Fgfr3* gene disrupts signaling pathways in cortical bone, we performed proteomics analyses of cortical bone from WT and *Hch* mice at 3,6,12 and 24 months of age (Extended Fig.4). In order to gain an understanding on the multivariate expression differences across ages and conditions (*Hch*), we performed weighted gene co-expression network analysis (WGCNA) and estimated the topological map of the proteins that demonstrated correlated expression patterns (Fig. 6a). We identified 13 clusters, with cluster 1, 5 and 10 capturing the highest percentage of the proteins showing correlated expressions (Fig. 6b and Extended Fig. 5a). Average cluster Z-score profiles across age and genotype revealed three broad trends: clusters that changed monotonically with age in both genotypes, clusters that diverged between WT and *Hch* at one or more ages, and clusters that combined both features (Fig. 6c).

**Figure 6:**
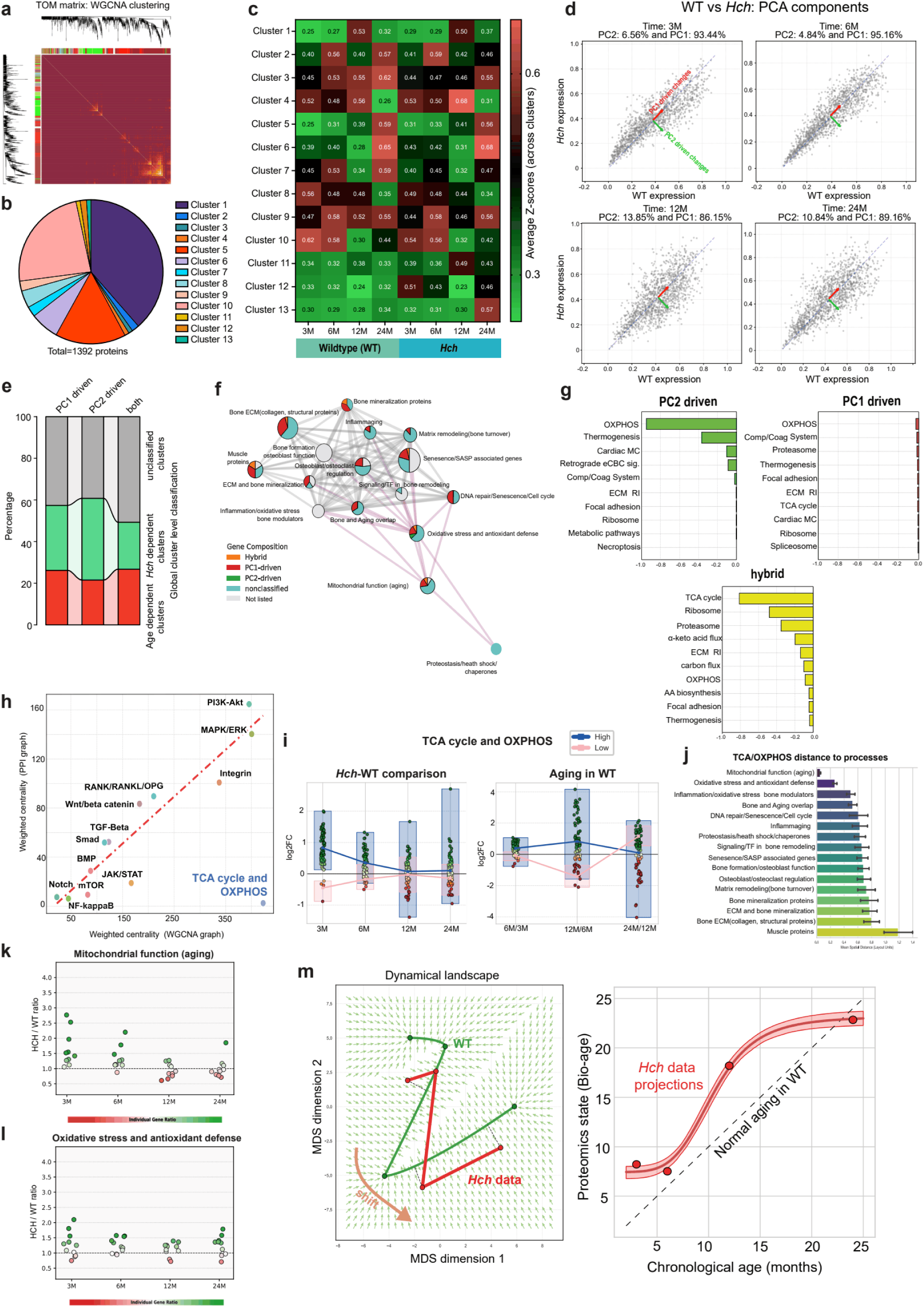
Mitochondrial dysfunction plays a key role in cortical bone of *Hch* mice. (**a**) Heatmap of the topological overlap matrix (TOM) for the gene network. In this heatmap, each row and column represent a gene. Lighter colors indicate lower topological overlap, whereas progressively darker red shades represent higher topological overlap. The darker blocks along the diagonal correspond to distinct gene modules. The gene dendrogram and the corresponding module assignments are displayed along the left and top margins of the heatmap. (**b**) Classification of the identified clusters based on their expression profiles. (**c**) Heatmap of normalized gene expression in identified gene clusters across WT and *Hch* samples at different ages. Rows represent gene clusters, and columns correspond to the experimental conditions (WT and *Hch* samples at different ages). Color intensity indicates normalized gene expression levels, ranging from low expression (green) to high expression (red). The numerical values within each cell represent the corresponding correlation coefficients. (**d**) *Hch* versus WT protein abundance at 3, 6, 12, and 24 months of age. Principal component analysis (PCA) of each point cloud separates age-specific deviations along the diagonal (PC1, 86–95% of variance) from genotype-specific deviation (PC2, 5–14%). Red and green arrows indicate the directions of age-driven and mutation-driven changes, respectively. (**e**)Classification of gene expression dynamics based on Principal Component (PC) contributions. Genes were classified into PC1 (Age-related), PC2 (Mutation-related), or Hybrid groups based on the mean percentage of variance explained across four time-integrated samples (**f**) Spatial network analysis. Significantly enriched biological processes and pathways are shown for each group. (**g**) Unbiased pathway enrichment analysis of the three datasets. (**h**) Comparative network analysis of pathway centrality across co-expression and physical interaction landscapes. Pathway centrality evaluation using both WGCNA, representing the actual experimental dataset, and Protein-Protein Interaction (PPI) graphs representing the underlying physical architecture. (**i**) Longitudinal expression dynamics (3,6,12 & 24-month-old mice) of TCA cycle and oxidative phosphorylation (OXPHOS) pathways (*Hch*-WT comparison and aging in WT respectively). (**j**) Proximity of the TCA/OXPHOS hub to downstream biological processes. (**k, l**) Longitudinal analysis (3,6,12 & 24-month-old mice) of mitochondrial function (aging) and oxidative stress and antioxidant defense pathways. (**m**) Multidimensional Scaling (MDS) where the wild-type (green) defines the natural aging “attractor” or path. Projection analysis to estimate “Bio-age” by projecting mutant data onto the WT trajectory.

In order to quantify these divergent trends, we performed a principal component analysis (PCA) across the Z-scores of the proteins at each age (Fig. 6d), and resolved the principal component (PC) contributions across *Hch* and WT. Across all four timepoints, PC1 captured 86–95% of the variance, indicating a time-dependent axis, whereas PC2 captured 5–14% of the variance, indicating contribution of the genotype to the expression variability (Fig. 6d). Furthermore, the protein abundance of *Hch* versus WT at each age across the PCs, demonstrated that the two components correspond to largely non-overlapping protein subsets whose relative contribution shifts with age (Fig. 6d).

Based on PC contributions, proteins were classified as age-related (PC1 contributions), *Hch* related (PC2 contribution), or hybrid, across four time-integrated samples (Fig. 6e and Extended Fig. 5b,c). The PC1shows high correlation between control and mutant Z-scores (PC1 > 90%), while PC2 explains >55% of the expression variance. Genes with PC contributions with 45%–55% explained variance were classified as hybrid, representing proteins with synergistic influence on both age and genotype (Fig. 6e). Unbiased Gene Ontology enrichment of the PC1-driven, PC2-driven, and hybrid protein demonstrates that 114 were shared across all groups, while 147 were unique to PC1 group, 46 to PC2 group, and 223 to hybrid, emphasizing that the hybrid class represents a distinct, convergent response rather than a simple sum of the two individual axes (Extended Fig. 5c). Based on these detailed classifications, a protein–protein interaction (PPI) network analysis followed by enrichment suggested segregation and functional interactions for the age-dependent and *Hch*-dependent clusters (Fig. 6f and extended Fig. 5d,e)

Together, these multivariate analyses reveal that *Fgfr3* gain-of-function alters cortical bone homeostasis through a pronounced genotype-by-age interaction that widens across the lifespan. By decoupling variance into distinct time-dependent and genotype-dependent axes, we show that canonical aging and *Hch*-driven features rely on largely non-overlapping biological machinery. Finally, the enrichment of unique pathways within the hybrid protein class demonstrates that *Fgfr3* hyperactivation synergizes with advancing age to generate an emergent, molecular phenotype rather than simply mimicking premature aging.

### Mitochondria and mitochondrial related processes are highly expressed in *Hch* mice cortical bone

PC1 exhibits a broad, “low-specificity” profile. While it shows high statistical enrichment, its influence is distributed across a wide array of pathways rather than a single functional unit. This suggests that aging acts as a systemic driver of rather than a precise “targetter” of a specific process (Fig. 6g). In contrast, the PC2 (*Hch* variant-specific) and Hybrid (*Hch* variant and age-specific) regimes demonstrate functional precision (Fig. 6g). Interestingly, we observed that these sets are highly targeted, with convergence on core metabolic hubs, specifically Oxidative Phosphorylation and the TCA Cycle. Analysis of *Hch* variant-driven proteins revealed a prominent mitochondrial signature, with eight main proteins (Ndufc2, Cox4i1, Acadvl, Slc25a4, Prdx3, Ssbp1, Gatd3, Fkbp10) involved in mitochondrial and oxidative phosphorylation were upregulated (Supplementary Excel Table). These data suggested high and primary involvement of mitochondria and mitochondrial related processes in *Hch* cortical bone (Figure 6g).

### The shift in TCA cycle and OXPHOS pathways is driven primarily by FGFR3 overactivation or compensatory effects

To determine the functional impact of the mutation, we evaluated pathway centrality using both WGCNA and Protein-Protein Interaction (PPI) graphs representing the underlying physical architecture (Fig. 6h and Extended Fig. 5f). First, we checked how strongly different signaling pathways contribute to PC2 regime and we saw that Wnt/β-catenin is a dominant driver of the *Hch* variant and other pathways such as Notch, BMP, JAK/STAT, RANK/RANKL/OPG, MAPK/ERK, and NF-κB highly contribute to the PC2 regime (Extended Fig. 5f). Next, we compared our network against the PPI network, which shows that most pathways follow the biological expectations where functional activity matches known physical structure. Interestingly, the TCA cycle and OXPHOS pathways exhibit significantly higher centrality in the WGCNA-based graph than in the PPI-based graph (Central in co-expression, but peripheral in PPIs) (Fig. 6h). This pattern indicates a gain-of-function or compensatory overactivity where the *Hch* variant forces that pathway to become a central coordination hub for the rest of the network. Our data suggests that the shift is driven primarily by overactivation or compensatory effects rather than a complete rewiring of the interactome. This indicates that the *Hch* variant is pushing existing metabolic machinery to an extreme functional state to maintain network coordination (Fig. 6h).

### The *Hch* mutation causes a premature boost in mitochondrial-related processes

To pinpoint the divergence between the experimental groups, we analyzed the *Hch*/WT Ratio (Fold Change) to observe the immediate impact of the mutation on the longitudinal expression dynamics of TCA cycle and OXPHOS pathways, at each stage (Fig. 6i). The results reveal a clear temporal acceleration in the HCH line. In pathways such as Oxidative Phosphorylation, *Hch* mice exhibit a high ratio relative to the WT at the earliest stages at 3 months, partial decrease at 6–12 months, and then appears to normalize at 24 months. Interestingly, the WT data shows a similar surge in activity, but only at the midpoint (12M/6 M) of the timeline before decreasing in the final step. This suggests that the mutation (*Hch* variant) causes a premature “spike” in these metabolic processes, forcing the mutant to reach an aged state much earlier than the WT, which only approaches that same state mid-process (Fig. 6i). This metabolic shift is functionally coupled to the PI3K-AKT and MAPK/ERK signaling axes, which rank among the most central pathways in both the PPI and WGCNA networks (Fig.6i). Longitudinal analysis revealed that the TCA/OXPHOS, MAPK/ERK and PI3K-Akt pathways follow broadly concordant trajectories (Extended Fig. 6,7) (other pathways are presented in Extended Fig. 6,7 as well). Additionally, we observed that mitochondrial function (aging) and oxidative stress and antioxidant defense processes exhibit the shortest network distances, identifying them as the primary “first responders” to metabolic shifts in the TCA cycle (Fig. 6j). Longitudinal analysis confirms that these primary interactors exhibit early-onset shifts in mutant-to-WT ratios at 3 months of age, supporting the conclusion that the TCA cycle initiates a cascading response through mitochondrial aging and oxidative stress pathways (Fig.6k,l). In conclusion, *Hch* variant reprograms the mitochondrial metabolism in bone cells.

### The aging trajectory of *Hch* mice differs from that of control mice

Finally, to get an understanding of the aging trajectories of the gene co-expression, we performed a multidimensional scaling (MDS). We observed that the *Hch* variant (red) causes a significant spatial shift in the landscape compared to the wild-type (green) that defines the natural aging (Fig. 6m). By projecting mutant data onto the control trajectory, we estimated “Bio-age” (Fig. 6m). While normal aging follows the diagonal (dashed line), the mutant exhibited a steep non-linear pattern above the main diagonal, which confirms that the mutation imposes an “aging burden” that saturates early. The mutant maintains an advanced biological age much earlier than the control (Fig. 6m). Both MDS and projection analysis reveal a “shifted” aging trajectory in the *Hch* cortical bone.

## Discussion

Hypochondroplasia is a well-characterized skeletal dysplasia caused by FGFR3 gain-of-function mutations ^1^, with prior studies largely focused on development and skeletal morphology ^26,27^. Patients are at increased risk of OA and its associated symptoms (joint pain, back pain, spinal cord compression), as well as hypotonia (muscle weakening) ^26^. To date, the interaction between *Hch* variants and musculoskeletal aging has not been studied. FGFR3 is predominantly expressed in chondrocytes, making it a central regulator of endochondral ossification and cartilage homeostasis during growth ^28,4^. Our findings show that FGFR3 also affects the musculoskeletal system during aging in the *Hch* mouse model ^6^. Aging is the most prominent risk factor for OA, aged chondrocytes characterized by telomere shortening, mitochondrial dysfunction, and elevated oxidative stress, resulting in a reduced capacity for matrix repair ^29^.

Few studies have considered a link between OA and FGFR3. One previous study found that *Fgfr3*^ach^ mice did not develop the typical bone thickening or joint damage seen during OA ^30^. In this present study, we observed increased subchondral bone remodeling in the medial compartment of the knee, suggesting OA-like changes. This divergence may arise from inherent difference in *Fgfr3* expression in different cell types. For instance, the *Fgfr3* gene is expressed ubiquitously in *Hch* mice ^6^ whereas in *Fgfr3*^ach^ mice ^31^, its expression is restricted to chondrocytes. The reduced articular cartilage thickness and abnormal thin collagen fibers in the medial tibial plateau of *Hch* mice could partially explain matrix metalloproteinase upregulation and extracellular matrix loss generally observed in OA ^32,33^. Our data suggest that FGFR3 variants increase the risk of articular cartilage degeneration.

Until now, the impact of FGFR3 mutations on muscle has not been directly investigated. It is well accepted that the interaction between bone and muscle extends beyond mechanical loading, with both tissues functioning as secretory organs that regulate each other through soluble factors, myokines and osteokines ^34^. Aging is accompanied by changes in the secretome of both bone and muscle, with a shift from beneficial, protective myokines (irisin, IL-15, and IGF-1) toward catabolic factors, pro-inflammatory cytokines (TNF-α and IL-6), leading to inflammaging that is now recognized as a central driver of sarcopenia ^35–37^. The muscle phenotype exhibited by *Hch* mice indicates alterations in myofiber size, processes supported by these bone-derived signals. The muscle weakness observed in our *Hch* model could therefore not only result from a disruption in bone signaling but also accelerate bone deterioration by reducing the myokines that maintain bone cell viability; a virtuous cycle that may underlie the concurrent progression of osteoporosis and sarcopenia ^38,25^. In Fgfr3-related skeletal dysplasia, the prevalence of ligamentous laxity is higher and associated with musculoskeletal problems ^39^. We hypothesize the presence of a direct link between ligamentous laxity and muscle abnormalities such as hypotonia, which could be further exacerbated by both *Fgfr3* variant and aging.

Bone metabolism is altered in ACH and HCH patients ^40^, as evidenced by reduced bone density; this has been confirmed in various *Fgfr3* mouse models ^41,42,6^. In this study, we found abnormalities in bone structure in the *Hch* model as aging progresses. μCT analysis of cortical bone revealed a dual effect. On one hand, an age-related decline in structural bone parameters is observed between 12 and 24 months in both genotypes. This is consistent with the well-characterized pattern of cortical bone aging in mice ^43–45^. On the other hand, a genotype-related increase in cortical BMD and BV/TV was observed in the *Hch* mice which indicates that increased FGFR3 signaling promotes cortical bone formation and/or suppresses cortical bone resorption. This finding is consistent with computed tomography analyses of the long bone performed in a cohort of ACH patients ^46^.

Interestingly, in spite of the differences in femoral length and bone structural parameters between the *Hch* and WT animals, matrix mineralization did not differ at 6 months of age and the same shift towards higher bone matrix mineralization was observed at 24 months of age in both genotypes indicating an impaired endochondral bone formation with no intrinsic mineralization defect. This is similar to what was seen in a mouse model of Hutchinson-Gilford Progeria Syndrome (HGPS), a fatal premature-aging disorder, in which qBEI revealed that bone matrix mineralization density distribution was statistically normal at all evaluated bone sites at both 8 and 15 weeks ^47^.

A notable finding was the presence of high intracortical porosity in some bone samples from aged mice of both genotypes, a feature that was absent from all samples obtained from 6-month-old mice. De novo intracortical remodeling associated with heterogeneous mineral density has previously been described as a characteristic feature of aged female mice ^48,44^. Our findings extend these observations by demonstrating comparable de novo intracortical remodeling in male mice. These morphological changes resulted in high intracortical porosity with osteon-like structures and new bone formation on the endosteal surfaces that was typically much less mineralized than the compact cortical shell of older non-remodeled bone tissue.

In our study, osteocyte canalicular density was decreased with age in both genotypes, in accordance with previous studies ^48^. We also measured an increase in osteocyte lacunar density in *Hch* mice; however, we do not know how many of these lacunae contained viable osteocytes. Our results show that the *Hch* gene variant had a significant impact on the size and number of osteocytes lacunae. It has previously been reported that osteocyte lacunae density decreases with age ^48^ or is unchanged ^44^, in contrast to our results. Note that these previous studies were mainly performed on female mice whereas our study was done on male mice.

The μCT analyses reported in this paper showed that bone quality was impaired in long bone leading to an increased risk of associated bone fragility. To identify the molecular changes underlying the observed microarchitectural phenotype, we performed a proteomic analysis of femoral cortical bone in mice from 3 to 24 months of age. Interestingly, the analysis of Fgfr3-variant-driven proteins revealed a prominent mitochondrial signature, with the main proteins (Ndufc2, Cox4i1, Acadvl, Slc25a4) seen to be involved in mitochondrial and oxidative phosphorylation. These mitochondrial pathways appeared earlier in time in *Hch* mice compared to WT, indicating that the mutation drives a premature acceleration of mitochondrial metabolism in cortical bone. Our data suggest that early mitochondrial overactivation in mutants represents the metabolic precursor to the secretory aging phenotype, in which senescent osteoblasts acquire a senescence-associated secretory phenotype (SASP) and release proinflammatory and ECM-degrading factors (Liang et al., 2024). Consistent with this, we detected inflammaging markers in PC2 (Il1rap, Siglec1, S100a11). Elevated OXPHOS activity increases ROS production, which can activate NF-κB-driven inflammatory signaling, ultimately promoting the osteoblast senescence and SASP central to age-related bone deterioration.

Moreover, assessing mitochondrial metabolism in *Hch* mouse articular cartilage, not only in bone, could be informative. Indeed, impaired mitochondrial metabolism in chondrocytes—altered OXPHOS, elevated oxidative stress, and failed mitophagy—drives cartilage degeneration, chondrocyte senescence, and OA progression (Qi et al., 2024; Guo et al., 2025; Danet et al., 2026).

Previous studies identified aging-related osteokines (AOK) by comparing young and aged C57BL6 male mouse bone and aging reprograms the bone secretome toward a SASP (senescence-associated secretory phenotype) ^49^. In the age-driven category of our data, Fgf23, Sost, and Acp5 were among the identified proteins and they fall within the BOK (Bone-specific OsteoKine) category as bone-enriched secreted proteins. ^49^. Sparc/osteonectin was also detected and it falls within the AOK category, showing decreased expression in aged bone and also appears in the myokine compilation functioning as a muscle-derived adipogenic inhibitor, placing it at the bone–muscle interface ^25,49^. Tgfbi overlaps both MOK (Mechanical-responsive OsteoKine) and OBK (OB-secreted OsteoKine), consistent with its dual regulation by mechanical stimuli and osteoblast secretion.

Several musculoskeletal proteins were also identified in this list, that are mainly contractile/structural proteins, ECM/collagen proteins and OXPHOS components such as Sod2. It is important to note that Sod2 was significantly upregulated at 3 months in *Hch* mice which reflects the cell’s attempt to neutralize the excess mitochondrial superoxide. However, SOD2 does not eliminate reactive oxygen species, it converts one (O2-) into another (H2O2). The production arm of the antioxidant cascade is activated, but the clearance arm fails to respond proportionally which leads to H2O2 accumulation within the mitochondrial matrix and cytosol of cortical bone cells.

Overall, our results show that aging in *Hch* mice is accompanied by a dual degenerative bone phenotype, combining impairment of muscle homeostasis and bone fragility in the long bones with OA-like changes in the articular cartilage and subchondral bone. Proteomic analysis further highlights mitochondrial dysfunction as a central mechanistic feature of this process, suggesting that bone mitochondria contribute to the loss of bone quality during aging and may therefore represent a promising therapeutic target for counteracting age-related musculoskeletal degeneration.

## Methods

### Mouse model

The *Fgfr3^Asn534Lys/+^* (*Hch*) mouse model was generated and characterized previously by Loisay et al.2023. Mice were maintained under specific pathogen–free conditions and used in accordance with institutional animal care guidelines. Genotyping was performed as previously described ^6^.

### Study approval

All procedures were approved by the French Animal Care and Use Committee (APAFIS #61235).

### Morphometric analyses

The samples consist of mouse male femurs (WT: 12-month-old *n* = 15, 24-month-old *n* = 11, *Hch*: 12-month-old *n* = 14, 24-month-old *n* = 12), and vertebrae (WT: 12-month-old *n* = 12, 24-month-old *n* = 11, *Hch:* 12-month-old *n* = 10, 24-month-old *n* = 11). μCT imaging of the femur and vertebrae was performed using Skyscan 1272 (Bruker). Samples were fixed in 70% ethanol and scanned with settings of 70 kV, 142 μA, 0.5 mm aluminium filter. 3D reconstructions were generated using NRecon software (Skyscan). In femurs, cortical bone parameters were determined at 5 μm voxel resolution of an ROI equal to 6% of the femur length centered in the mid-shaft bone 59% along the proximo-distal length of the femur, and trabecular parameters were obtained at 5 μm voxel resolution in an ROI equal to 6% of the femur length in size beginning 0.6% proximal to the distal growth plate. Trabecular bone parameters of L5 vertebrae 5 were determined at 5 μm voxel resolution by analyzing the whole vertebral body region delimited by the 2 cartilage endplates. Trabecular bone parameters of subchondral bone were determined at 5 μm voxel resolution by analyzing the subchondral bone region delimited by the subchondral bone plate and the growth plate.

### Bone samples preparation

Femurs were obtained from 6-month-old and 24-month-old male mice. For each age group, WT mice (n = 10) and *Hch* mice (n = 10) were analyzed. The femurs were fixed in 70% ethanol and transversely sectioned. Proximal and distal bone segments were then dehydrated through ascending ethanol concentrations, defatted in acetone, and embedded in polymethylmethacrylate (PMMA) according to standard procedures ^18,47^. During the embedding procedure, the proximal segment was stained with rhodamine to allow for the subsequent visualization of the osteocyte lacuno-canalicular network (OLCN) ^50,51^. The staining was performed by using a mixture of ethanol and rhodamine (1.25 g rhodamine per 300 ml 100% ethanol). Samples were placed in tubes containing staining fluid (14 ml) for 3 days, with daily renewal of the staining fluid, on a homemade low speed spinning wheel, leading to a complete penetration with rhodamine of the sample ^50,51^.

After hardening, all samples were shaped into cuboidal blocks using a diamond saw. The longitudinal section of the distal part of the femurs were exposed by cutting using a hard tissue microtome (SM2500, Leica, Nussloch, Germany), whereas cross-sections at the midshaft were prepared for the proximal, rhodamine-stained samples. Thereafter, all samples were ground and polished using a polishing device (PM5 Logitech, Glasgow, UK).

### Quantitative backscattered electron imaging (qBEI)

Sample preparation and qBEI measurements are described in detail elsewhere ^18,47,20,52^. Briefly, the longitudinal sections of the distal femur were carbon coated using vacuum evaporation (Agar SEM Carbon coater, Agar Scientific Limited, Essex, UK) to ensure electric conductivity of the surface. Backscattered electron images were collected using a field emission scanning electron microscope (FESEM, SEM SUPRA 40, Zeiss) at 20 kV, 10 mm working distance and 270–320 pA sample current with 0.88 μm spatial resolution. A high purity aluminum/carbon reference sample was used for calibration to compare between measurements ^53^.

#### a) Bone Mineralization Density Distribution (BMDD)

The metaphyseal spongiosa (ms), epiphyseal (es) and cortical (cort) sites of distal femurs were analyzed separately. The qBEI images of the murine femurs were processed and evaluated using in-house written ImageJ (https://imagej.nih.gov/ij/) macros to provide the grey level histogram that was further converted to weight percent calcium (weight % Ca). From the quantitative backscattered electron microscopy image showing the mineral content in each pixel, the corresponding frequency distribution, the bone mineralization density distribution (BMDD) is extracted (note that areas of lowly mineralized, highly porous cortical bone tissue were excluded from qBEI measurements).

The BMDD histogram is characterized by five parameters ^53^.

- CaMean (weight % Ca); the average calcium concentration,
- CaPeak (weight % Ca); the most frequently measured calcium concentration,
- CaWidth (Δ weight % Ca); the full width at half maximum of the curve, characterizing the heterogeneity in matrix mineralization,
- CaLow (% bone area); the percentage of lowly mineralized bone, defined as the area which is mineralized below the 5^th^ percentile of the BMDDs of the 6 months old WT animals for each femoral region as follow: 16.12 weight % Ca (ms); 19.24 weight % Ca (es) and 24.09 weight % Ca (cort).
- CaHigh (% bone area); the percentage of highly mineralized bone area, which is mineralized above the 95th percentile the BMDDs from the 6 months old WT separately for each femoral region as follow: 29.29 weight % Ca (ms); 29.46 weight % Ca (es) and 29.81 weight % Ca (cort).

### b) Osteocyte lacunae section (OLS) characteristics

The osteocyte lacunae sections (OLS) of the femurs were analyzed on 6 qBEI images of the longitudinal sections covering on average 0.9 ± 0.2 mm² mineralized bone area containing an average of 438 ±94 lacunae. The analysis of osteocyte lacunae sections was performed using a custom-made macro in ImageJ (https://imagej.nih.gov/ij/). The images were binarized using a threshold of 5.2 weight % Ca to discriminate mineralized bone matrix from OLS. The OLS were further filtered using a size threshold between 3 µm^2^ and 80 µm^2^ as described previously for the analysis of murine bone ^20^.

The OLS characteristics are described by six parameters.

- OLS-density (OLS number/mm^2^); defined as the number of OLS per bone area,
- OLS-porosity (%); defined as the OLS total area per bone area,
- OLS-area (µm^2^); median value of the OLS area per sample,
- OLS-perimeter (µm); median value of the OLS perimeter per sample,
- OLS aspect-ratio: the median value of the OLS aspect ratio of each sample. The aspect ratio is given by the ratio of long to short axis of a fitted ellipse to the OLS and is a measure of anisotropy of the OLS, where a value of 1 denotes a circle, and elongation along one axis of the pore results in an aspect ratio of >1. Aspect ratio values >10 were excluded from the analyses.
- OLS nearest neighbor distance (NND, µm), the median distance of the nearest centroid to each OLS.

### Characterization of the Osteocyte Lacuno-canalicular Network (OLCN)

The osteocyte lacuno-canalicular network of rhodamine-stained samples (n=7, for each group) was visualized by confocal laser scanning microscopy (CLSM, TCS SP5, Leica, Wetzlar, Germany). An oil-immersion objective (HCXPL APO 63x/140-60 OIL) with an airy 1 pinhole of 95.5 µm was used. The measurements were done on four regions of interest (ROI). For each region we obtained stacks of images with 1024×1024 pixels with lateral resolution 0.24 µm/pixel and a step size of 0.25 µm in z-direction reaching to a depth of approximately 60 µm in the sample. The excitation wavelength of the used He-Ne Laser was 543 nm. The laser intensity and the photomultiplier gain voltage were continuously adjusted in depth to guarantee sufficient image quality. The images were analyzed using the TINA software package developed by F. Repp and co-workers ^50,51^. TINA is a collection of Python scripts to binarize, skeletonize and analyze the obtained images. The main result of the analysis is the osteocyte lacuno-canalicular density measured as total canalicular length per volume and expressed as μm/μm^3^.

### Knee joint histology

Whole leg was collected in 4% paraformaldehyde, fully decalcified in 0.5 M EDTA (pH 7.5), and paraffin embedded. Serial sections (8 μm) were used for Safranin O or Sirius red staining. Images were captured with an Olympus PD70-IX2-UCB microscope.

### RNA extraction

Total RNA was extracted from tibialis anterior muscles of 12-month-old male (WT *n* = 12, *Hch n* =13). mice using TRIzol Reagent (Invitrogen, Thermo Fisher Scientific) according to the manufacturer’s instructions. Briefly, the muscles were homogenized in TRIzol, followed by phase separation with chloroform. RNA was precipitated from the aqueous phase with isopropanol and stored overnight at −80°C. The next day, the samples were centrifuged, washed with 75% ethanol the next day, resuspended in RNase-free water and then stored at −80°C.

### RT-qPCR

Complementary DNA (cDNA) was synthesized from 100-1000 ng of total RNA using the SuperScript IV Reverse Transcriptase kit (Thermo Fisher Scientific) with random hexamer or oligo(dT) primers, following the manufacturer’s protocol. Quantitative PCR was performed using HOT FIREPol® EvaGreen® qPCR Supermix (Solis BioDyne) on a thermocycler. Cycling conditions consisted of an initial denaturation at 95°C for 12 minutes, followed by 42 cycles of 95°C for 15 seconds, 59°C for 20 seconds, and 72°C for 20 seconds, with a final melt curve analysis to confirm amplicon specificity. All reactions were performed in technical triplicates. Relative gene expression (Trim63 and MyoD) was calculated using the 2^−ΔΔCt method, normalized to the reference gene (RpL7L1).

### Immunofluorescence

Tibialis anterior muscle tissues were sectioned transversely at 10 µm using a cryostat and mounted onto SuperFrost Plus slides (12-month-old: WT *n* = 6, *Hch n* =6). Sections were allowed to air-dry at room temperature for 30 minutes, then fixed in 4% paraformaldehyde in phosphate-buffered saline (PBS) for 10 minutes at room temperature, followed by three washes in PBS. Sections were blocked with 5% bovine serum albumin (BSA) in PBS for 30 minutes at room temperature in a humidified chamber. Sections were incubated with wheat germ agglutinin conjugated to Alexa Fluor 488 (WGA–AF488; Thermo Fisher Scientific, 1:1000) overnight at 4°C in the dark. Slides were then washed three times in PBS and nuclei were counterstained with DAPI. Sections were mounted with aqueous fluorescence mounting medium and imaged on a confocal microscope. Muscle fiber cross-sectional area (CSA) and Shape Factor Index (SFI) were quantified from WGA-stained images using FIJI/ImageJ.

### Grip Strength Test

Forelimb grip strength was assessed in 3 and 12-month-old male mice (WT *n* = 6, n=19 respectively, *Hch n* =6, *n* =12 respectively) using a grip strength meter (ugo basile). Each mouse was held by the tail and allowed to grasp a horizontal metal grid with its forelimbs only. The animal was then pulled horizontally away from the grid at a steady pace until it released its grip, and the peak force (g) was recorded. Each mouse performed two trials with a few minutes rest interval between trials, and the average of the two measurements was used for analysis. All tests were performed at the same time of day to minimize circadian variability.

### Proteomics sample preparation

Femurs from 3, 6, 12 & 24-month-old male mice (WT n = 7, n=7, n=7 & n=3, *Hch* n = 7, n=7, n=7 & n=3 respectively) were dissected and stripped of surrounding muscle tissue. Epiphyses were removed, and bone marrow was eliminated by centrifugation to obtain clean cortical bone tissue and minimize contamination from non-osseous components. Bones were demineralized by incubation in 1.2 M hydrochloric acid (HCl, 1 mL per sample) with overnight rotation at 4°C. Prior to grinding, all homogenizer components, including 2.0 mL impact-resistant screw-cap tubes, homogenizing plates, and steel beads, were pre-cooled with liquid nitrogen. Individual frozen femurs were pulverized using a MM 400 mixer mill (Retsch) with steel grinding beads for 30 seconds at 20 Hz. Beads were subsequently removed and cleaned.

### Protein extraction

Proteins were extracted following the protocol described by ^54^. S-Trap™ 96-well plate S-TrapTM (Protifi, Hutington, USA) digestion was performed on 30µg of protein extract according to manufacturer’s instructions. Briefly, samples were supplemented with 20% SDS to a final concentration of 5%, reduced with 20mM TCEP (Tris(2-carboxyethyl) phosphine hydrochloride) and alkylated with 50mM CAA (chloracetamide) for 5min at 95°C. Aqueous phosphoric acid was then added to a final concentration of 2.5% following by the addition of S-Trap binding buffer (90% aqueous methanol, 100mM TEAB, pH7.1). Mixtures were then loaded on S-Trap plate. Five washes were performed for thorough SDS elimination. Samples were digested with 1,5 µg of trypsin (Promega) at 47°C for 2h. After elution, peptides were vacuum dried and resuspended in 2% ACN, 0.1% formic acid (FA) in HPLC-grade water. 600ng were injected on an Evosep One system coupled to a timsTOF HT (Bruker Daltonics, Germany) mass spectrometer. The Evosep One system operated with the Whisper Zoom 40

Samples Per Day method using a 15 cm C18 Aurora Elite column (AUR3-15075C18-CSI, IonOpticks). The mobile phases were 0.1% FA in H_2_O (solvent A) and 0.1% FA in ACN (solvent B). Mass-spectrometric data were acquired using the parallel accumulation serial fragmentation (PASEF) acquisition method in DIA mode. diaPASEF setting were: mass range from 475 to 1000 m/z, mobility ranges from 0.85 to 1.27 V s/cm^2^ (1/k0), cycle time estimate of 0.95s, mass width of 25Da and 21 mass steps for cycle.

### Weighted Gene Co-expression Network Analysis (WGCNA)

Weighted Gene Co-expression Network Analysis (WGCNA) provides a powerful framework for uncovering these hidden patterns in gene or protein expression data (<u>WGCNA: an R package</u> <u>for weighted correlation network analysis | BMC Bioinformatics | Springer Nature Link</u>). WGCNA takes into account similar protein expression patterns across multiple samples. This co-expression often indicates functional relationships and creates weighted correlations, indicating that the continuous nature of co-expression relationships.

Using WGCNA, we computed a correlation coefficient matrix between every pair of genes across all samples (WT and *Hch*, age agnostic). All ages of the experimental samples were considered. Missing values were removed from analysis if >50% of the values were absent across each sample-age replicate. This creates a similarity matrix where high values indicate strong co-expression. The correlation values are raised to a power (β), which emphasizes strong correlations while suppressing weak ones. We used a soft threshold of 12, creating a weighted adjacency matrix of the expression patterns. Typically, WGCNA uses the topological overlap measure (TOM) as proximity. A TOM was calculated measuring not just direct correlation between two genes, but also how many neighbors they share. Using the TOM dissimilarity (1-TOM) as a distance measure, we performed hierarchical clustering to group genes/proteins into modules or clusters. 13 modules/clusters were identified. A dynamic tree-cutting algorithm identifies these modules automatically, with the dynamic tree cut threshold fixed at 0.65. For each module or cluster, WGCNA calculates a “module eigengene” (ME), essentially the first principal component of the module’s expression matrix.

### Identification of super clusters from WGCNA modules

To comprehensively map the molecular landscape of the murine models, we systematically evaluated and compared the higher-order network organization across a multi-factorial framework encompassing genotype (WT vs. *Hch*) and chronological age (3M, 6M, 12M, and 24M) using the Z-scores obtained from WGCNA. These Z-scores reveal whether the cooperative relationships between genes remain stable or are actively rewired and fluctuating with respect to baseline controlsBy tracking the global Z-score profile across the 13 identified modules, we decoupled and identified the independent contributions of age and mutation separately within the dataset, specifically isolating the distinct impact of aging within the mutant condition.

### Gene set enrichment analysis

To perform functional enrichment on the genes in Age-dependent clusters and *Hch*-dependent clusters, an over-representation analysis based on the hypergeometric distribution was conducted across three distinct databases such as Kyoto Encyclopedia of Genes and Genomes (KEGG) database (<u>KEGG Database</u>), Reactome (<u>Home - Reactome Pathway Database</u>) and Gene Ontology (GO) datasets (<u>Gene Ontology Resource</u>), applying a Benjamini-Hochberg adjusted p-value and q-value cutoff of 0.05. Gene Ontology (GO) enrichment evaluated biological processes, cellular components, and molecular functions. The organism was set for Mus musculus.

### Protein-protein interaction network

The protein-protein interaction (PPI) functionality from Metascape (http://www.metascape.org) was used to generate PPI maps of proteins specific to mutations and age. Metascape relies on integrating and scoring physical PPI pairs from eight resources (BIND, BioGRID, DIP, IntAct, MatrixDB, NetPath, Reactome, WikiPathways), as the proxy of all public-domain physical PPI data sources. Metascape applies Molecular Complex Detection (MCODE) algorithm (<u>An automated method for finding molecular complexes in</u> <u>large protein interaction networks - PMC</u>) to identify tightly connected network cores.

### Gene-space principal component partitioning (GPCP) for the separation of aging, genotype, and combined expression effects

To systematically decouple the individual and combined impacts of chronological age and genetic mutation on the proteome, we developed a gene-space principal component partitioning (GPCP) framework. We used the average Z-scores per month for both WT and *Hch* conditions across the time series (3M to 24M).

A row-wise Z-score normalization was performed which ensures that every protein contributes equally to the directional model based on the shape of its profile rather than its min-max compressed magnitude.

Following this, for every time step (3M, 6M, 12M, 24M), the protein expression levels were mapped onto a two-dimensional coordinate system where the X-axis represents the WT expression values WT and the Y-axis represents the *Hch* expression levels. Principal Component Analysis (PCA) for each time step as a dimensionality reducing method to identify the proteins contributing to the most variance. The first PCA axis aligns closely with the main diagonal, capturing the shared, correlated transcriptomic variance, while the the second PCA axis runs delineates the discordant, non-correlated variance. The resulting PCA coordinates were converted into a variance-driven ratio (*PC_i_*^2^/(*PC_i_*^2^ + *PC_j_*^2^)), allowing genes to be explicitly categorized by whether their variance was primarily dictated by shared baseline biology (PC1) or mutant-specific divergence (PC2). Subsequently, for both the WT and mutant conditions, the time-mean (3-24months) of the variance-driven ratio for both principal components was calculated. Genes were categorized into three distinct functional subgroups based on their final relative variance scores:

PC1 group: Genes driven by PC1, defined by a PC1 variance-driven ratio> 0.9
PC2 group: Genes driven by PC2, defined by a PC2 variance-driven ratio≥0.55.
Hybrid group: A hybrid group influenced by the coordinates of both axes, where both PC1 and PC2 variance-driven ratios fell between 0.45 and 0.55.

The expression of genes falling into the Hybrid group was likely shaped by a combination of both physiological aging and mutation-driven effects.

### Process-based network construction and visualization

Based on the groups (PC1, PC2 and hybrid driven) we focused on several different processes and their associated genes/proteins involved within the context of the mutation (Supplementary methods - Biological process_list sheet). Based on this, the PPI network of the genes/proteins involved in the processes was built based on STRING database high confidence score 0.7 [<u>STRING: functional protein association networks</u>]. From the PPI network of the genes/proteins involved within the listed processes, we calculated the Topological Overlap Matrix (TOM). An edge was constructed between two processes if their mutual mean TOM was greater than zero.

For each valid interaction, the edge weight was defined directly by the mean TOM value. Concurrently, the topological distance between the nodes was calculated as 1 - mean TOM value. A distance approaching 0 indicates that the two processes possess an almost identical gene/protein overlap. Network visualization was performed using the NetworkX library via a force-directed spring layout (nx.spring_layout). Furthermore, the internal distribution of genes/proteins across PC1, PC2, and hybrid regimes was mapped directly onto the respective node distributions.

### Unbiased pathway enrichment and topology-based alteration analysis

To mitigate potential bias arising from imbalances in dataset sizes, specifically, the proportionally larger size of the PC1 gene set (378 genes/proteins) relative to the PC2 and Hybrid sets (51 and 71 genes/proteins respectively), a targeted network expansion was applied to the PC2 and Hybrid gene/protein sets. To expand the candidate gene lists for the PC2 and Hybrid regimes, we identified first-degree neighbors within the STRING protein-protein interaction (PPI) network that overlapped with the differentially expressed proteins (DEPs). Interacting partners were retained only if they maintained a minimum of two shared connections with the core experimental dataset. Interactions were filtered using a stringent confidence score threshold of 0.7. Through this expansion, the PC2 and Hybrid gene/protein sets were increased to 147 and 302 candidates, respectively [Supplementary methods - pc2_pc1_hybrid_core, PC2 expansion, hybrid expansion sheets].

Functional pathway enrichment analysis was performed across all three regimes and their associated genes/proteins (PC1, PC2, and Hybrid). For each statistically enriched pathway, an enrichment signal score was extracted from the STRING database, with higher scores signifying a more robust biological enrichment.

Initial screening of enriched pathways across the three sets, revealed substantial topological and biological redundancy within the Hybrid regime [Supplementary methods - PC1_PC2_hybrid_enriched pathways].

Subsequent enrichment scores were stratified and retained according to their derivation from the PC1, PC2, or Hybrid datasets.

The operational baseline was systematically and computationally depleted of PC1-, PC2-, and Hybrid-specific genes, respectively, to measure the residual impact of each set on the topology of the baseline enriched pathways. Network topological measures were evaluated across the complete enriched pathway graphs to quantify functional centrality. The absolute change in network connectivity of a specific pathway graph was subsequently quantified by calculating the change in degree centrality pre- and post-computational deletion of the specific genes/proteins identified in the PC1, PC2, and Hybrid datasets. A pronounced decrease in network connectivity, yielding a negative degree change, demonstrated that the deleted experimental protein group functions as a critical structural hub or central pillar within that specific pathway network, meaning their removal destabilizes the underlying pathway integrity.

A composite Impact Score was formulated as a multiplication between the degree change and enrichment signal score associated. By multiplying the topological measurement by the statistical confidence score, the final Impact Score selectively prioritizes pathways that are both simultaneously overrepresented in the data and structurally dependent on the specific gene/protein group identified in this study.

### Pathway interaction modeling

#### Selection of target pathways

To conduct pathway interaction analysis, we selected a list of target pathways using both a candidate-driven and an unbiased approach. Gene sets for these selected pathways were defined by taking the union of the corresponding processes annotated in the KEGG and Reactome databases, provided that these genes or proteins were identified in the experimental dataset.

#### Construction of a pathway interaction graph based on experimental data analysis

To capture the functional relationships between biological pathways mediated by co-expression networks, a pathway-pathway interaction network was constructed using Weighted Gene Co-expression Network Analysis (WGCNA) modules.

To identify pathway co-occurrences within identical functional modules, the integrated dataset was subjected to an inner self-join operation on the cluster identifier key (ClusterID). This operation generated an exhaustive pairwise matrix of all pathways that shared membership within the same WGCNA cluster. Identical pathway pairs mapping to themselves (self-loops) were filtered out of the resulting matrix. The edge weight (strength) between any two pathways was defined as the total number of distinct WGCNA clusters they mutually shared. To identify the core biological pathways driving the network architecture, a normalized weighted degree centrality metric was calculated for each node.

#### Construction of a pathway interaction graph based on the PPI network

To model the functional interactions and cross-talk between biological pathways, a pathway-level graph was constructed using protein-protein interaction (PPI) data as a baseline scaffolding. The reference PPI network was retrieved from the STRING database, applying a high-confidence threshold (combined score 0.7). Pathway-pathway edges were established based on the presence of shared PPIs. These edges were weighted by the total frequency of inter-pathway protein interactions to represent cross-talk density. Ultimately, normalized weighted degree centrality was computed for each node to highlight the key biological pathways governing the network structure

#### Comparative analysis of expression-based and physical interaction networks

We compared the node centralities of the dataset-specific WGCNA graph against the database-derived PPI graph to identify condition-specific network rewiring. Pathways aligning with the diagonal line represent a baseline state where functional expression matches the known physical architecture. Deviations from this diagonal indicate distinct biological phenomena: a shift toward the WGCNA axis denotes a functional gain-of-function or compensatory overactivity. Conversely, a shift toward the PPI axis indicates a functional deficit, where the physical blueprint of the pathway is present but transcriptionally inactive.

Spatial network analysis and proximity of the TCA/OXPHOS hub to downstream biological processes.

The mean spatial distance was calculated based on a network overlay across multiple iterations to identify which biological processes are most strongly coupled with the TCA/OXPHOS metabolic hub. In this overlay, the proximity of nodes is determined by gene-set overlap, with processes sharing a high number of genes being positioned closer together.

### Quantification of transcriptomic state-space trajectories and aging dynamics

To characterize the temporal proteomics trajectories, Multidimensional Scaling (MDS) was performed on the gene expression profiles across wild-type (WT) and mutant cohorts at chronological intervals of 3, 6, 12, and 24 months. MDS was selected to preserve the global relative distances between high-dimensional profiles in a lower-dimensional space. Dissimilarity matrices were constructed using pairwise Euclidean distances across all timepoints, and the high-dimensional data matrix was projected into a two-dimensional space using an automated normalized stress minimization criterion

To model a continuous baseline of healthy physiological aging, a smooth WT reference manifold was generated by fitting a Piecewise Cubic Hermit Interpolating Polynomial (PCHIP) across the empirical WT coordinate matrices over the time domain in (3, 24) months. The PCHIP formulation was specifically selected to preserve local monotonicity and prevent numerical overshoots or non-biological oscillations between discrete empirical timepoints.

Mutant trajectories were mapped relative to the continuous WT reference manifold to determine biological transcriptomic age. For each discrete mutant data point the corresponding equivalent biological age and structural path drift were derived by identifying the orthogonal projection or minimal Euclidean distance to the fine-grained WT manifold

To model the transition dynamics of the mutant transcriptomic state toward an altered homeostasis, a constrained Hill equation or “soft switch” sigmoidal function was fitted to the derived biological ages. To regularize the fitting procedure against overfitting given discrete sampling, parameter search bounds were strictly enforced. Statistically robust variance and 95% confidence intervals (CI) for the predicted non-linear trajectories were determined via a bootstrapping framework (N = 200), where in empirical data points were iteratively perturbed with Gaussian noise (μ = 0, σ= 0.4) prior to objective function minimization. All computational pipelines, numerical interpolations, and curve fits were executed utilizing NumPy, SciPy, and Scikit-Learn libraries.

### Statistics

Statistical analyses were performed using GraphPad Prism version 10 (GraphPad Software, San Diego, CA, USA). Two-way ANOVA was used to assess the effects of two independent variables and their interaction. Groups were compared in a nonparametric Mann–Whitney *U* test or Student’s *t* test as appropriate. Results were considered statistically significant at p < 0.05. Data are presented as mean ± standard deviation (SD).

## Acknowledgments

We thank the Imagine Institute’s imaging facility and the Structure Fédérative de Recherche’s histology facility for their help with this work. We thank Amélie de La Seiglière, Nabil Kaci, Léa Loisay, Emilie Dambroise and Emilie Panafieu (LEAT-Imagine Institute-SFR Necker) for providing critical advice during the course of study. CF, GG and MT were funded by the European Union within the Horizon Europe MSCA program under agreement No 101072766 (CHANGE Project). TYCHE software was used for articular cartilage analysis. This project received a grant from the National Research Agency (Bony Brain: ANR-23-CE17-0030). We would like to thank the philanthropy department of Mutuelles Axa, through the “Head and Heart Chair,” for supporting the Imagine institute’s imaging facility. We thank the « Fond de dotation Regain » for financial support. This study was supported by the Austrian Social Health Insurance Fund (OEGK), the Austrian Workers’ Compensation Board (AUVA). The authors thank Dr. Jochen Zwerina, Director of the Ludwig Boltzmann Institute of Osteology, Vienna for continuous support; Petra Keplinger, Sonja Lueger, and Phaedra Messmer for careful qBEI measurements. SB, MAH and NFZ and are part of the European Reference Network on Rare Bone Diseases (www.ernbond.eu).

## Author contributions

CF designed and performed the experiments and LLM directed the research project. GG, SB, MT, MH, AF, SC, IM performed some experiments. GG, SB, MT, MH, ICG, PB, FZ, NFZ analyzed the results and CF made the figures. DM and SS analyzed the proteomics results. NFZ, FZ, PB CF and LLM wrote the manuscript. All authors reviewed and critically edited the manuscript.

