## extended figures and tables for "Mitochondrial dysfunction plays a key role in premature musculoskeletal aging in hypochondroplasia mouse model"

#### Extended Figure 1: Trabecular bone is affected in *Hch* mice during aging

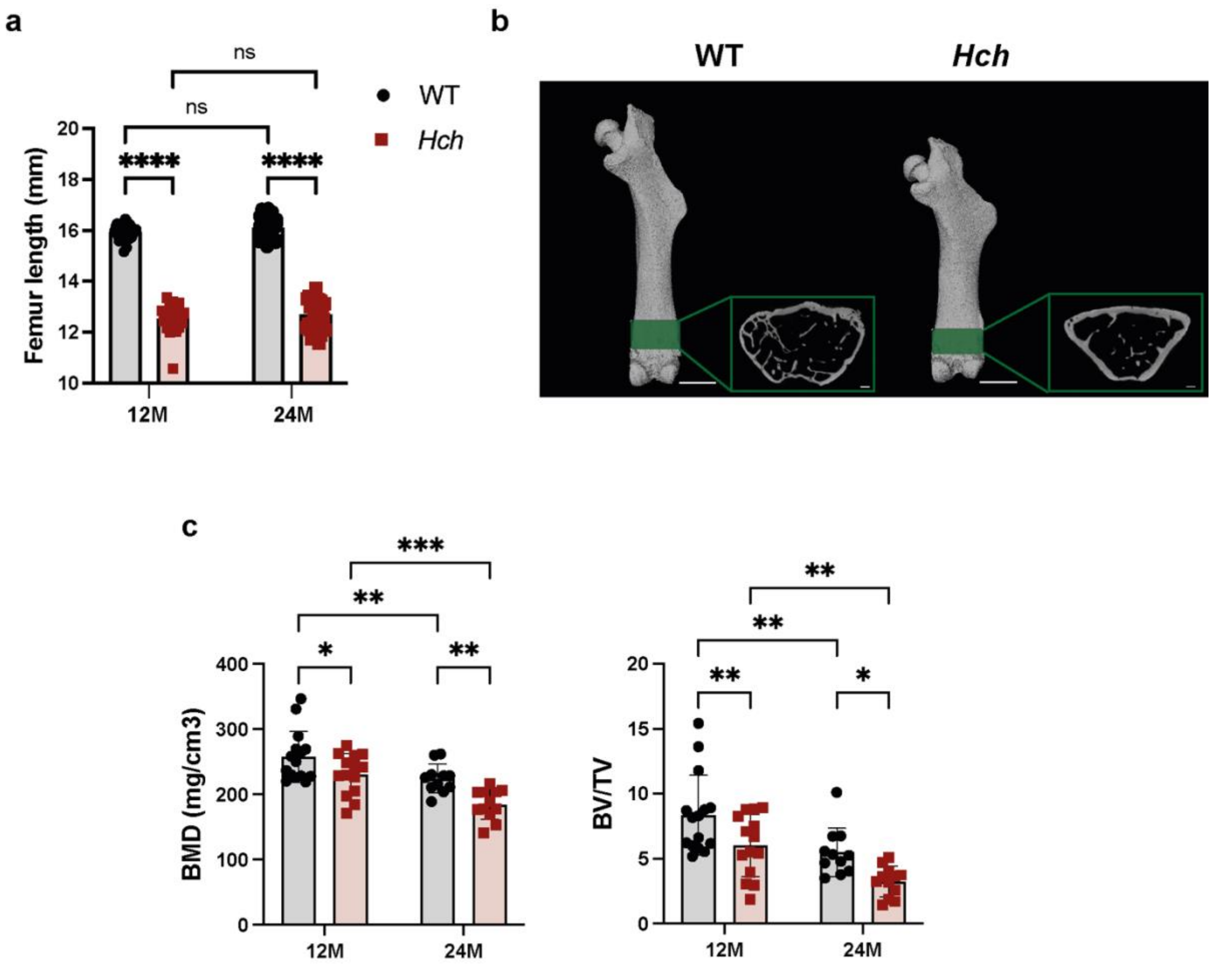


(**a**) Histograms representation of femurs' length at 12 and 24 months of age in *WT* ($n$ = 37; n=29) vs *Hch* ($n$ = 24; n=35). (**b**) 3D visualization of representative 12-month-old *WT* and *Hch* male femurs. Highlighted in green indicates the ROI chosen for the trabecular bone studies. Scale bars: 500 µm for the whole bone and Scale bars: 200 µm for trabecular parts. (**c**) Histograms for the trabecular bone parameters of 12 & 24-month-old femurs of *Hch* mice (n=15, n=11 respectively) vs WT mice (n=14, n=12 respectively): Bone mineral density (BMD) and ratio of bone volume to tissue volume (BV/TV). Data are presented as mean ± SD. (****, $P < 0.0001$; ***, $P < 0.001$; **, $P < 0.01$; *, $P < 0.05$; ns, non-significant; Mann-Whitney for 2 groups and two-way ANOVA for >2 groups).

**Extended Figure 2:** Bone mineralization and Ca content are normal in *Hch* mice

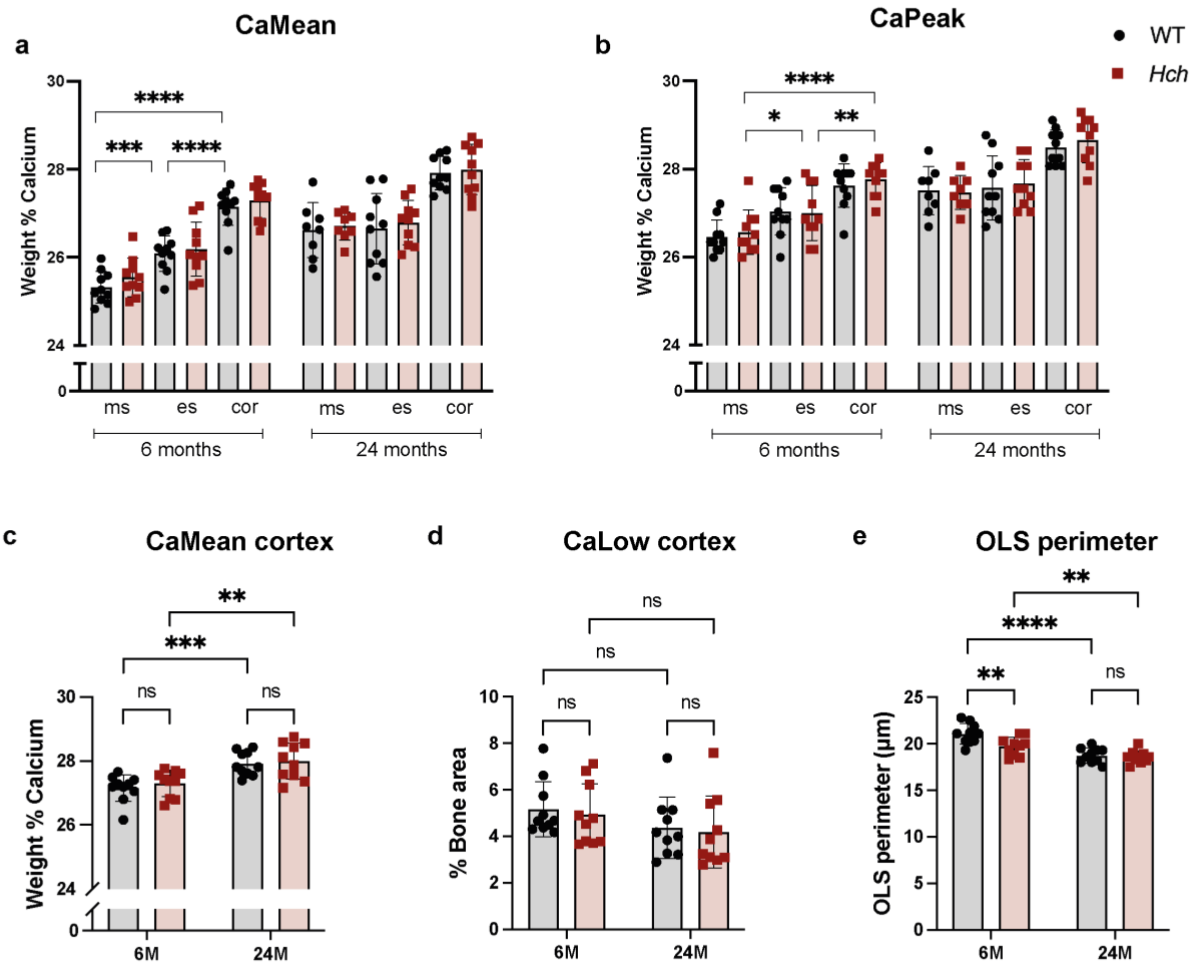


(**a**) Statistical comparison of CaMean and (**b**) CaPeak of *Hch* mice (n=10) vs WT mice (n=10) in the cortex, epiphysis, and metaphysis. (**c**) Statistical comparison of CaMean and (**d**) CaLow of *Hch* mice (n=10) vs WT mice (n=10) in the cortex. (**e**) Statistical comparison of OLS perimeter in *Hch* (n = 10) versus WT mice (n = 10) mice femoral cortex. CaMean: the average calcium concentration. CaPeak: the most frequently measured calcium concentration. CaLow: the percentage of lowly mineralized bone. Data are presented as mean ± SD. (****, $P < 0.0001$; ***, $P < 0.001$; **, $P < 0.01$; *, $P < 0.05$; ns, non-significant; Mann-Whitney for 2 groups and two-way ANOVA for >2 groups).

#### Extended Figure 3: OA scoring and collagen fibers distribution

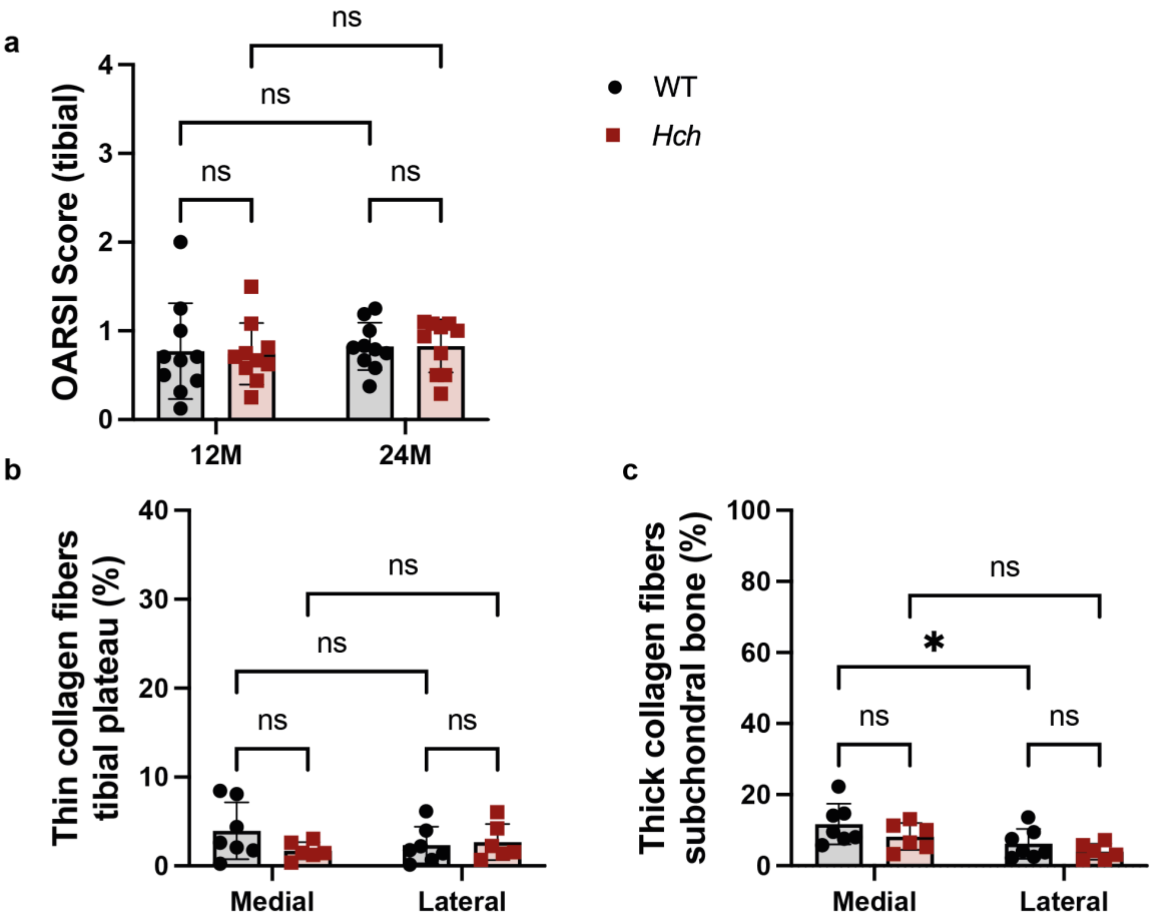


(**a**) OARSI score for 12 and 24-month-old WT (n=10 for both) and *Hch* mice (n=10 for both). (**b**) Percentage of thin collagen fibers in medial and lateral tibial plateaus and (**c**) percentage of thick collagen fibers in medial and lateral parts of the subchondral bone of 12-month-old mice (*WT n* = 7; *Hch n* = 6). Data are presented as mean ± SD. (****, $P < 0.0001$; ***, $P < 0.001$; **, $P < 0.01$; *, $P < 0.05$; ns, non-significant; Mann-Whitney for 2 groups and two-way ANOVA for >2 groups).

#### Extended Figure 4: Proteomic protocol

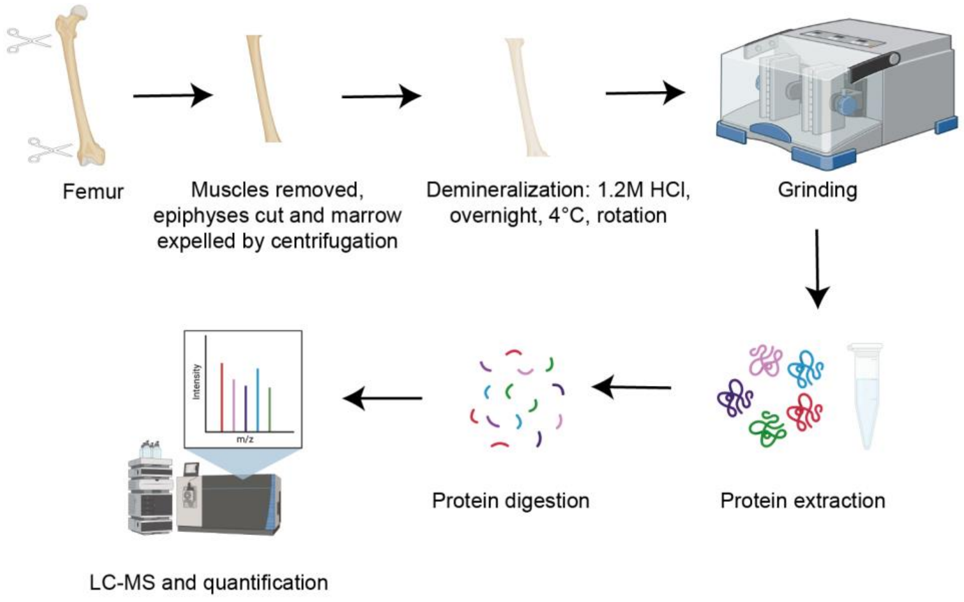


Design of the proteomic protocol for cortical bone

#### Extended Figure 5: Clustering and PPI networks

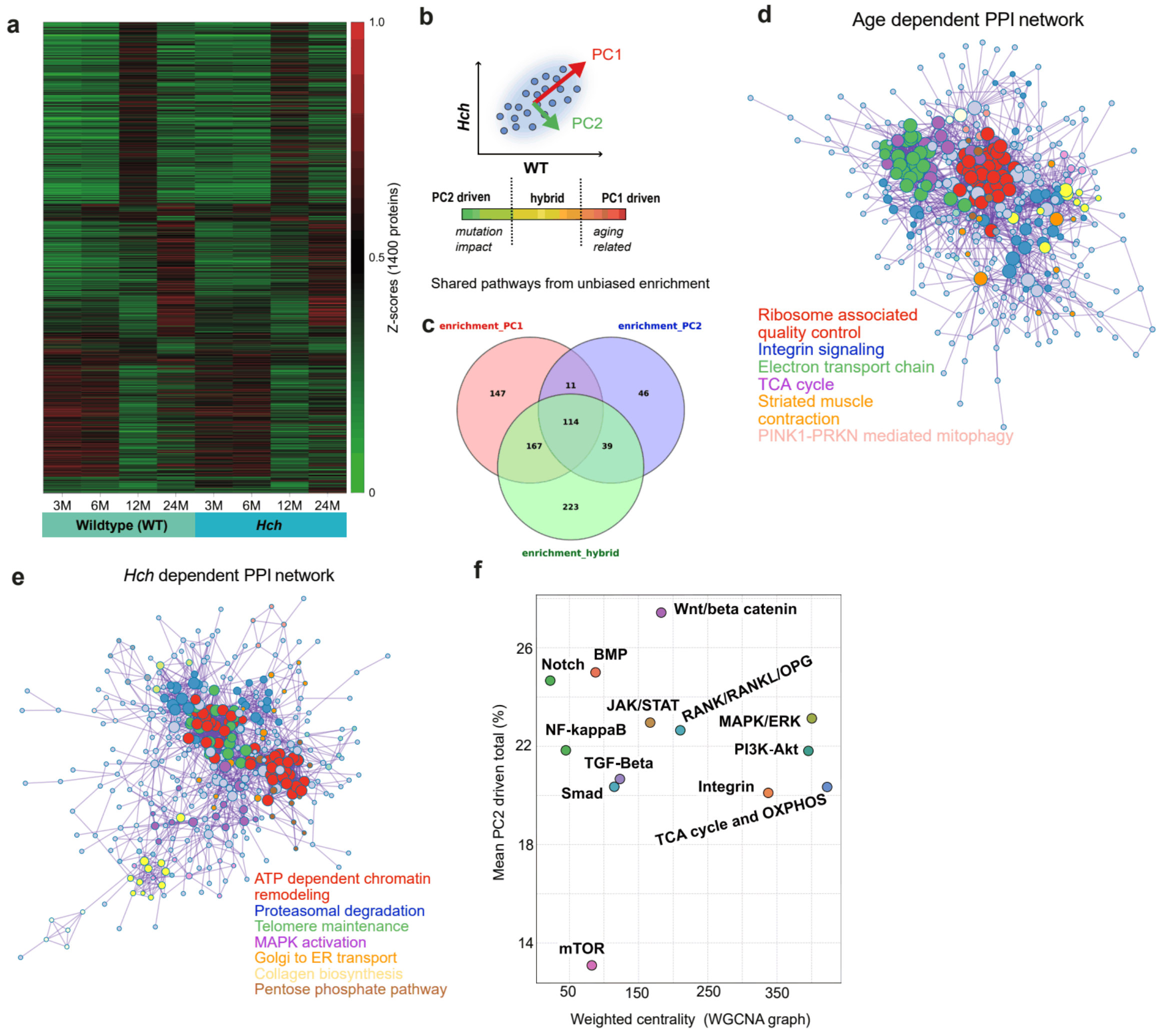


(**a**) Z-score normalized heatmap displaying differential expression patterns between *WT* and *Hch* groups across time. Row scaling highlights relative expression shifts, identifying distinct patterns of expression. (**b**) Classification of gene expression dynamics based on Principal Component (PC) contributions. Genes were classified into PC1 (Age-related), PC2 (Mutation-related), or Hybrid groups. (**c**) Gene set enrichment analysis of the age and *Hch* dependent clusters using Gene Ontology (GO) and/or KEGG pathway annotations. Significantly enriched biological processes and pathways are shown for each group. (**d,e**) Age dependent and HCH dependent Protein–protein interaction (PPI) network of the identified proteins. Nodes represent individual proteins and are colored based on their pathway assignments. Edges indicate protein– protein interactions, illustrating the functional relationships among the proteins. (**f**) Weighted Gene Co-expression Network Analysis (WGCNA).

### Extended Figure 6: Changes in various pathways in WT during aging

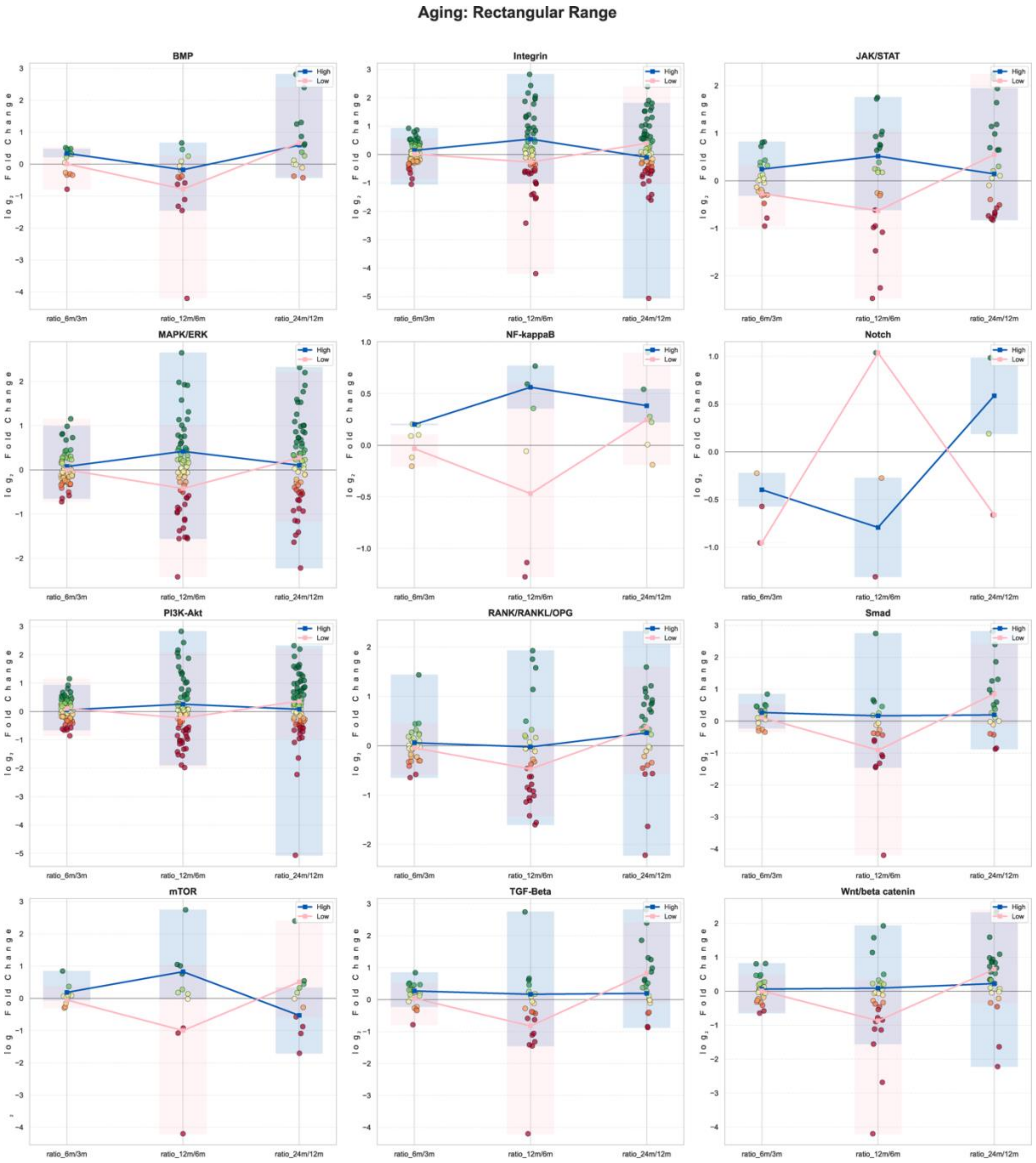


Longitudinal expression dynamics of different pathways in WT male mice with aging.

#### Extended Figure 7: Changes in various pathway in Hch vs WT

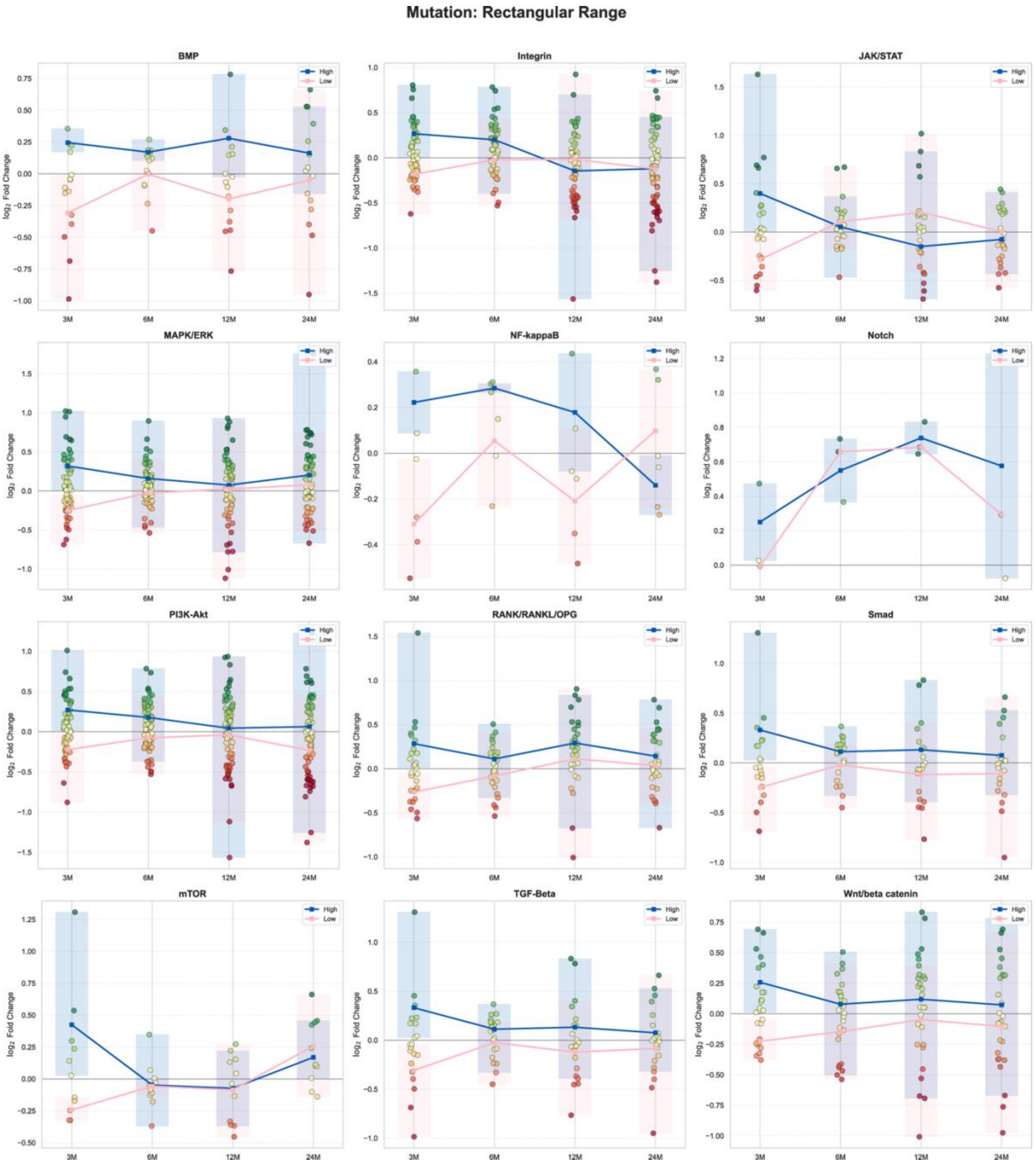


Longitudinal expression dynamics of different pathways in *Hch* vs WT (y axis represents HCH/WT) at 4 timepoints (x axis: 3,6,12 and 24 months of age).

### Supplementary Tables

#### Supplementary Table 1

Femur length measurements of both male and female mice at 12 & 24 months. 2way Anova test, values presented as mean ± standard deviation.

| Femur Length | Mean values (± SD) | | | | 2-Way ANOVA | | |
|---|---|---|---|---|---|---|---|
| | WT 12 months | *Hch* 12 months | WT 24 months | *Hch* 24 months | Effect of genotype | Effect of age | Interaction term |
| Femur length (mm) | 15,933 (0.297) | 12,524 (0.571) | 16,108 (0.451) | 12,677 (0.568) | <0.0001 | 0.0594 | 0.9008 |

#### Supplementary Table 2

Femur trabecular measurements obtained with µCT are presented and classified by genotype and age. Values are expressed as mean ± standard deviation. P-values of the 2-Way ANOVA are reported.

| Trabecular bone parameters | Mean values (± SD) | | | | 2-Way ANOVA | | |
|---|---|---|---|---|---|---|---|
| | WT 12 months | *Hch* 12 months | WT 24 months | *Hch* 24 months | Effect of genotype | Effect of age | Interaction term |
| BMD (mg/cm$^3$) | 257.944 (39.272) | 231.321 (31.372) | 223.405 (21.985) | 184.884 (22.951) | 0.0003 | <0.0001 | 0.4548 |
| BV/TV | 8.352 (3.077) | 6.018 (2.383) | 5.504 (1.861) | 3.242 (1.160) | 0.0008 | <0.0001 | 0.9560 |

#### Supplementary Table 3

Femur cortical measurements obtained with µCT are presented and classified by genotype and age. Values are expressed as mean ± standard deviation. P-values of the 2-Way ANOVA are reported.

| Cortical bone parameters | Mean values (± SD) | | | | 2-Way ANOVA | | |
|---|---|---|---|---|---|---|---|
| | WT 12 months | *Hch* 12 months | WT 24 months | *Hch* 24 months | Effect of genotype | Effect of age | Interaction term |
| TMD ($mg/cm^3$) | 1616.225 (22.550) | 1658.815 (41.848) | 1603.765 (79.062) | 1613.465 (47.639) | 0.0645 | 0.0418 | 0.2398 |
| BV/TV | 41.065 (2.135) | 45.413 (3.257) | 37.742 (2.135) | 37.076 (4.709) | 0.0079 | <0.0001 | 0.1476 |

#### Supplementary Table 4

Comparison of CaMean and CaPeak at different bone sites. One-Way ANOVA, repeated measures. ms: metaphyseal spongiosa, ep: epiphysis, cort: cortical bone

| | P-value | | |
|---|---|---|---|
| | ms vs. ep | ms vs. cort | ep vs. cort |
| CaMean: WT 6 months | 0.0009 | <0.0001 | <0.0001 |
| CaMean *Hch* 6 months | 0.0048 | <0.0001 | 0.0002 |
| CaMean WT 24 months | 0.3709 | 0.0005 | <0.0001 |
| CaMean *Hch* 24 months | 0.4043 | 0.0001 | <0.0001 |
| CaPeak: WT 6 months | 0.0033 | <0.0001 | 0.0024 |
| CaPeak *Hch* 6 months | 0.0209 | <0.0001 | 0.0020 |
| CaPeak WT 24 months | 0.2696 | 0.0009 | 0.0004 |
| CaPeak *Hch* 24 months | 0.1141 | <0.0001 | <0.0001 |

#### Supplementary Table 5

Femur metaphyseal spongiosa BMDD parameters obtained with qBEI are presented and classified by genotype and age. Values are expressed as mean ± standard deviation. P-values of the 2-Way ANOVA are reported.

| Metaphyseal spongiosa (ms) | | | | | | | |
|---|---|---|---|---|---|---|---|
| BMDD parameters | Mean values (± SD) | | | | 2-Way ANOVA | | |
| | WT 6 months | *Hch* 6 months | WT 24 months | *Hch* 24 months | Effect of genotype | Effect of age | Interaction term |
| CaMean (weight % Ca) | 25.32 (0.37) | 25.54 (0.44) | 26.62 (0.63) | 26.71 (0.32) | 0.2910 | <0.0001 | 0.6767 |
| CaPeak (weight % Ca) | 26.46 (0.38) | 26.57 (0.51) | 27.51 (0.55) | 27.47 (0.38) | 0.8428 | <0.0001 | 0.6313 |
| CaWidth (Δ weight %Ca) | 3.57 (0.54) | 3.172 (0.32) | 3.465 (0.31) | 3.661 (0.33) | 0.4476 | 0.1536 | 0.0306 |
| CaLow (% bone area) | 4.96 (0.89) | 4.89 (0.68) | 4.42 (0.83) | 4.23 (0.46) | 0.5902 | 0.0217 | 0.8160 |
| CaHigh (% bone area) | 4.750 (2.50) | 6.19 (3.30) | 18.04 (8.42) | 18.43 (2.84) | 0.5659 | <0.0001 | 0.7391 |

#### Supplementary Table 6

Femur epiphysis BMDD parameters obtained with qBEI are presented and classified by genotype and age. Values are expressed as mean ± standard deviation. P-values of the 2-Way ANOVA are reported.

| Epiphysis (es) | | | | | | | |
|---|---|---|---|---|---|---|---|
| BMDD parameters | Mean values (± SD) | | | | 2-Way ANOVA | | |
| | WT 6 months | *Hch* 6 months | WT 24 months | *Hch* 24 months | Effect of genotype | Effect of age | Interaction term |
| CaMean (weight % Ca) | 26.09 (0.41) | 26.19 (0.61) | 26.66 (0.80) | 26.79 (0.50) | 0.5349 | 0.0036 | 0.9229 |
| CaPeak (weight % Ca) | 27.03 (0.54) | 27.00 (0.63) | 27.57 (0.73) | 27.67 (0.54) | 0.8544 | 0.0035 | 0.7304 |
| CaWidth (Δ weight %Ca) | 2.96 (0.29) | 2.77 (0.14) | 3.69 (0.51) | 3.59 (0.38) | 0.2261 | <0.0001 | 0.7140 |
| CaLow (% bone area) | 4.99 (0.90) | 4.92 (0.58) | 4.85 (1.15) | 5.13 (0.54) | 0.6889 | 0.9007 | 0.5191 |
| CaHigh (% bone area) | 5.41 (2.76) | 5.97 (4.05) | 13.58 (9.42) | 15.27 (7.56) | 0.5881 | 0.0001 | 0.7854 |

#### Supplementary Table 7

Femur cortical BMDD parameters obtained with qBEI are presented and classified by genotype and age. Values are expressed as mean ± standard deviation. P-values of the 2-Way ANOVA are reported.

| Corticalis (cort) | | | | | | | |
|---|---|---|---|---|---|---|---|
| BMDD parameters | Mean values (± SD) | | | | 2-Way ANOVA | | |
| | WT 6 months | *Hch* 6 months | WT 24 months | *Hch* 24 months | Effect of genotype | Effect of age | Interaction term |
| CaMean (weight % Ca) | 27.15 (0.42) | 27.30 (0.41) | 27.91 (0.38) | 28.00 (0.57) | 0.4245 | <0.0001 | 0.8234 |
| CaPeak diaphysis (weight % Ca) | 27.62 (0.49) | 27.76 (0.38) | 28.49 (0.39) | 28.66 (0.52) | 0.2808 | <0.00001 | 0.9002 |
| CaWidth (Δ weight %Ca) | 3.03 (0.19) | 3.09 (0.19) | 3.05 (0.19) | 3.09 (0.31) | 0.4524 | 0.8015 | 0.9828 |
| CaLow (% bone area) | 5.17 (1.18) | 4.93 (1.33) | 4.37 (1.31) | 4.19 (1.55) | 0.6211 | 0.0802 | 0.9444 |
| CaHigh (% bone area) | 5.37 (2.90) | 6.41 (3.11) | 14.62 (6.47) | 18.85 (9.96) | 0.1942 | <0.0001 | 0.4298 |

#### Supplementary Table 8

Femur cortical OLS parameters are presented and classified by genotype and age. Values are expressed as mean ± standard deviation. P-values of the 2-Way ANOVA are reported.

| OLS parameters | Mean values (± SD) | | | | 2-Way ANOVA | | |
|---|---|---|---|---|---|---|---|
| | WT 6 months | *Hch* 6 months | WT 24 months | *Hch* 24 months | Effect of genotype | Effect of age | Interaction term |
| Porosity (%) | 0.8050 (0.12) | 0.819 (0.05) | 0.7280 (0.08) | 0.760 (0.084) | 0.7389 | 0.01656 | 0.3964 |
| Density (nb/mm$^2$) | 425.4 (42.2) | 473.7 (33.7) | 487.9 (42.24) | 513.6 (50.18) | 0.0092 | 0.0005 | 0.4066 |
| Area (µm$^2$) | 17.16 (1.07) | 15.82 (0.94) | 13.58 (0.92) | 13.82 (0.60) | 0.0601 | <0.0001 | 0.0084 |
| Perimeter (µm$^2$) | 21.07 (1.10) | 19.72 (0.99) | 18.74 (0.80) | 18.57 (0.67) | 0.0117 | <0.0001 | 0.0465 |
| Aspect-ratio | 4.15 (0.33) | 3.88 (0.43) | 4.21 (0.28) | 4.113 (0.29) | 0.0913 | 0.1823 | 0.4146 |
| NND (µm) | 26.80 (1.5) | 25.71 (0.73) | 25.58 (0.95) | 24.86 (0.98) | 0.0117 | 0.0044 | 0.5903 |

#### Supplementary Table 9

Femur OLCN parameter is presented and classified by genotype and age. Values are expressed as mean ± standard deviation. P-values of the 2-Way ANOVA are reported.

| OLCN Parameter | Mean values (± SD) | | | | 2-Way ANOVA | | |
|---|---|---|---|---|---|---|---|
| | WT 6 months | *Hch* 6 months | WT 24 months | *Hch* 24 months | Effect of genotype | Effect of age | Interaction term |
| OLCN density µm/ µm$^3$ | 0.33 (0.02) | 0.34 (0.03) | 0.25 (0.02) | 0.28 (0.02) | 0.0968 | <0.0001 | 0.3743 |

#### Supplementary Table 10

Lumbar vertebrae 5 measurements obtained with µCT are presented and classified by genotype and age. Values are expressed as mean ± standard deviation. P-values of the 2-Way ANOVA are reported.

| L5 measurements | Mean values (± SD) | | | | 2-Way ANOVA | | |
|---|---|---|---|---|---|---|---|
| | WT 12 months | *Hch* 12 months | WT 24 months | *Hch* 24 months | Effect of genotype | Effect of age | Interaction term |
| Vertebral height (mm) | 3.096 (0.113) | 2.846 (0.099) | 3.296 (0.151) | 3.041 (0.070) | <0.0001 | <0.0001 | 0.9378 |
| Interpedicular distance (mm) | 1.988 (0.049) | 1.893 (0.088) | 1.988 (0.041) | 1.967 (0.055) | 0.0022 | 0.0442 | 0.0442 |
| Spinal canal area ($mm^2$) | 1.178 (0.231) | 0.985 (0.174) | 1.206 (0.241) | 0.981 (0.072) | 0.0004 | 0.8220 | 0.7755 |

#### Supplementary Table 11

Lumbar vertebrae 5 measurements are presented and classified by genotype and age. Values are expressed as mean ± standard deviation. P-values of the 2-Way ANOVA are reported.

| Trabecular bone parameters (L5) | Mean values (± SD) | | | | 2-Way ANOVA | | |
|---|---|---|---|---|---|---|---|
| | WT 12 months | *Hch* 12 months | WT 24 months | *Hch* 24 months | Effect of genotype | Effect of age | Interaction term |
| BMD (mg/cm$^3$) | 428.454 (18.715) | 454.031 (21.778) | 369.665 (25.408) | 414.189 (38.595) | 0.0002 | <0.0001 | 0.2721 |
| BV/TV | 22.668 (1.648) | 24.644 (1.752) | 17.802 (2.176) | 20.753 (2.900) | 0.0004 | <0.0001 | 0.6745 |
| Tb.Nb (mm) | 3.889 (0.197) | 4.615 (0.365) | 3.228 (0.662) | 3.798 (0.572) | 0.0001 | <0.0001 | 0.6114 |

#### Supplementary Table 12

Articular cartilage measurements obtained with Safranin'O staining are presented and classified by genotype and age. Values are expressed as mean ± standard deviation. P-values of the 2-Way ANOVA are reported.

| Cartilage measurements | Mean values (± SD) | | | | 2-Way ANOVA | | |
|---|---|---|---|---|---|---|---|
| | WT 12 months | *Hch* 12 months | WT 24 months | *Hch* 24 months | Effect of genotype | Effect of age | Interaction term |
| Medial Cartilage Thickness (μm) | 101.000 (6.784) | 91.982 (9.252) | 99.228 (8.444) | 83.857 (8.799) | <0.0001 | 0.0712 | 0.2420 |
| Lateral Cartilage Thickness (μm) | 86.417 (9.589) | 77.045 (7.139) | 87.680 (4.148) | 78.946 (8.097) | 0.0005 | 0.5097 | 0.8940 |
| Medial Cartilage Calcfied Area (%) | 52.957 (3.100) | 57.315 (4.387) | 53.071 (5.487) | 52.044 (4.907) | 0.2553 | 0.0819 | 0.0698 |
| Lateral Cartilage Calcfied Area (%) | 46.328 (5.449) | 43.035 (8.225) | 42.577 (6.360) | 45.854 (7.937) | 0.9971 | 0.8365 | 0.1513 |

#### Supplementary Table 13

Articular cartilage measurements obtained with Picosirius red staining are presented and classified by genotype. The difference among WT and *Hch* mice is also presented in percentage and the relative p-value is reported.

| | | | 12 months | | | |
|---|---|---|---|---|---|---|
| | | | WT n=7 ; *Hch* n=6 | | | |
| | | Genotype | Mean | SD | Difference (%) | P value |
| Articular Cartilage Measurements | Thick collagen fibers MTP (%) | WT | 27.257 | 11.633 | -56,700% | 0.0013 |
| | | *Hch* | 11.802 | 4.515 | | |
| | Thick collagen fibers LTP (%) | WT | 15.573 | 5.785 | -27.171% | ns |
| | | *Hch* | 11.342 | 5.146 | | |
| | Thin collagen fibers Medial SB (%) | WT | 2.446 | 1.324 | +136.822% | 0.0093 |
| | | *Hch* | 5.792 | 3.184 | | |
| | Thin collagen fibers Lateral SB (%) | WT | 2.164 | 2.104 | +51,764% | ns |
| | | *Hch* | 3.284 | 1.424 | | |

#### Supplementary Table 14

Subchondral bone measurements obtained with µCT are presented and classified by genotype and age. Values are expressed as mean ± standard deviation. P-values of the 2-Way ANOVA are reported.

| Trabecular bone parameters (subchondral bone) | Mean values (± SD) | | | | 2-Way ANOVA | | |
|---|---|---|---|---|---|---|---|
| | WT 12 months | *Hch* 12 months | WT 24 months | *Hch* 24 months | Effect of genotype | Effect of age | Interaction term |
| BMD ($mg/cm^3$) | 539.500 (37.374) | 589.470 (53.615) | 486.40 (48.102) | 460.033 (35.610) | 0.3974 | <0.0001 | 0.0095 |
| BV/TV | 27.254 (2.719) | 31.070 (3.973) | 24.254 (3.074) | 22.198 (2.713) | 0.4385 | <0.0001 | 0.0071 |
| Tb.Nb (mm) | 3.842 (0.226) | 4.421 (0.389) | 3.214 (0.308) | 3.702 (0.295) | <0.0001 | <0.0001 | 0.6429 |

#### Supplementary Table 15

Peak muscle force values are presented and classified by genotype and age. Values are expressed as mean ± standard deviation. Values are expressed as mean ± standard deviation. P-values of the 2-Way ANOVA are reported.

| | | 12 months | | | |
|---|---|---|---|---|---|
| | | WT n=19 ; *Hch* n=12 | | | |
| GRIP test | Genotype | Mean | SD | Difference (%) | P value |
| Peak Force Value (gf) | WT | 91.028 | 39.851 | -54.766% | 0.0003 |
| | *Hch* | 41.175 | 14.385 | | |

#### Supplementary Table 16

Muscles' fibers analyses are presented and classified by genotype. Values are expressed as mean ± standard deviation. The difference among WT and *Hch* mice is also presented in percentage and the relative p-value is reported.

| | | | 12 months<br>WT n=6 ; *Hch* n=6 | | | |
|---|---|---|---|---|---|---|
| | | Genotype | Mean | SD | Difference (%) | P value |
| Muscles' Fibers Analyses | Cross-sectional area ($\mu m^2$) | WT | 2255 | 369.6 | -27.289% | 0.0087 |
| | | *Hch* | 1639 | 260.1 | | |
| | Minimal feret's diameter (µm) | WT | 44.71 | 4.205 | -17.442% | 0.0087 |
| | | *Hch* | 36.91 | 2.971 | | |
| | Shape Factor Index | WT | 1.423 | 0.021 | +1.997% | 0.0411 |
| | | *Hch* | 1.452 | 0.020 | | |

Link for Supplementary methods for proteomics: Supplementary Methods_Proteomics.xlsx
